# NEUROLINGUA: A Neuroimaging Database Tailored to Unravel the Complexity of Multilingual Comprehension

**DOI:** 10.1101/2025.06.06.658050

**Authors:** Ileana Quiñones, Amaia Carrión-Castillo, Iñigo Diez, Laura de Frutos-Sagastuy, Brendan Costello, David Carcedo, Lucía Manso-Ortega, Maksim Slivka, Abraham Sánchez, Anique Schüller, César Caballero-Gaudes, Pedro M. Paz Alonso, Manuel Carreiras

## Abstract

The neural mechanisms underlying language processing involve a well-defined brain network, including mainly left perisylvian areas. Yet, the extent of individual variability remains largely unexplored, particularly in bilingual and multilingual contexts. Differences in linguistic profiles (e.g., age of acquisition, exposure, proficiency) provide an opportunity to assess how network topology is shaped by sociolinguistic factors. To address this, we developed NEUROLINGUA, a comprehensive database of functional and structural MRI data, enriched with sociodemographic, sociolinguistic, and behavioral information. It includes 725 healthy individuals aged 18-82 immersed in a Basque-Spanish multilingual environment, ranging from near-monolinguals to highly proficient multilinguals. Participants completed a functional MRI language localizer with both auditory and visual comprehension tasks, enabling cross-modal comparisons, as well as sentences involving arithmetic problem-solving. Exploratory analyses confirmed associations between structural MRI, sociodemographic, and cognitive measures. We demonstrate that NEUROLINGUA’s functional MRI data localize the language comprehension network and thus capture linguistic profile effects. This integrative dataset offers an unparalleled resource to investigate factors influencing language network adaptability and variability in diverse sociolinguistic contexts.

## Background and Summary

Regardless of the structural diversity of human languages, their fundamental essence lies in the need to transmit information between individuals. Understanding the neural mechanisms underlying this process has been the focus of extensive research ^1–5^. Current neuroanatomical models describing the neural substrate of language comprehension propose that the linguistic system comprises a network of distributed gray matter regions, functionally connected via long- and short-distance white matter fiber tracts ^1–6^. Specifically, functional interactions between predominantly left-hemisphere areas—temporal, frontal, and parietal—are essential for both written and spoken language comprehension ^5,7^. The functional areas within this network have been delineated according to both their cytoarchitectonic features and functional properties ^8^.

Although previous neuroanatomical models ^1,3,9–11^ have provided key insights into the neurobiology of language comprehension, several questions remain unresolved. The bilingual and multilingual brain exemplifies these knowledge gaps, with still a limited understanding of the neural architecture underlying language comprehension in contexts where multiple languages coexist ^5,7,12–18^. This study aims to bridge this gap through NEUROLINGUA, a comprehensive database of functional and structural MRI data, enriched with sociodemographic, sociolinguistic, and behavioral information. This database has been built from data collected across multiple research projects conducted at the Basque Center on Cognition, Brain and Language (BCBL) over recent years ^7,12,17–24^.

NEUROLINGUA includes data from 725 healthy individuals, aged 18-82, with linguistic profiles ranging from near monolingual (of Spanish) to balanced bilinguals (of Spanish and Basque). This linguistic diversity enables in-depth investigation of the neural networks underlying language comprehension, including cross-linguistic differences and inter-individual variability linked to distinct linguistic profiles. All participants have been assessed with a functional MRI (fMRI) language localizer that includes both auditory and visual comprehension tasks ^25^, providing a relevant perspective for cross-modal comparisons. Additionally, this functional localizer includes experimental conditions in which auditory or visually presented sentences lead participants to perform simple mathematical operations. This added dimension is an advantage over existing databases, allowing researchers to identify the neurobiological basis of mental computation while controlling for associated linguistic processes.

The philosophical shift in the practice of science, driven by the open science movement, has encouraged many research groups to share data from language studies. This paper contributes to this initiative through NEUROLINGUA, which allows researchers to investigate the flexibility of the language network and its capacity for context-driven adaptation. By integrating NEUROLINGUA with existing databases ^26–28^, we will unlock aspects of the language networks previously inaccessible to study, marking a significant advance in result reproducibility—an improvement we hope will become standard practice. Unlike the existing databases, NEUROLINGUA enables exploration of the neurobiological underpinnings of language comprehension in bilingual and multilingual contexts. With NEUROLINGUA, we can address new questions in language comprehension research: Can the language network reorganize the functions of its critical nodes based on task demands? Do information flow and network topology vary according to sociolinguistic factors such as age of acquisition, exposure, and proficiency? Does this network flexibility depend on demographic factors like age and sex? What functional and structural plastic changes occur as a result of managing several languages?

## Methods

### Recruitment of the NEUROLINGUA cohort

To achieve a detailed characterization of the functional networks supporting language comprehension, while accounting for variability in participants’ linguistic profiles, we developed NEUROLINGUA, a harmonized database encompassing demographic, behavioral, and neuroimaging—functional and structural—data. Participants included in this database were recruited through calls launched by the BCBL between January 2010 and October 2022 for various projects. Recruitment efforts utilized informational leaflets, advertisements, and outreach on the BCBL website, institutional social media platforms, and even ads on local public transportation, ensuring a broad age range of participants. The BCBL also has a website where individuals interested in participating in research studies can register as users. Upon registration, they are assigned an anonymized code and, based on their profile, receive notifications about studies seeking participant profiles that match theirs. This website not only facilitates the management of participants and appointments but also features a questionnaire for potential participants. This questionnaire allows for a detailed characterization of each individual’s linguistic and socio-demographic profile.

Individuals who responded to these calls voluntarily took part in various research projects conducted during this period, all of which received prior approval from the BCBL Ethics Committee. Following the Declaration of Helsinki for research involving human subjects, all participants provided informed consent authorizing the use of their data for research purposes beyond a specific scientific question. This consent outlined the technical aspects of MRI, the purpose of the research, and the intended use of the collected data. It included permission to share data derived from the projects with investigators outside the BCBL. Additionally, each participant completed a safety questionnaire to identify any contraindications for MRI scanning. The BCBL Ethics Committee has approved the use of these data for the creation of a database under reference number 130525AT.

### Sample description and data harmonization

Given the variability in data collection over the years and the intrinsic changes in laboratory infrastructure, an extensive harmonization effort was undertaken. This process involved a detailed analysis of the collected variables and the definition of inclusion criteria:

- Participants had to be over 18 years of age, have normal or corrected-to-normal vision and hearing, and have no history of neurological, psychiatric, or language disorders.
- Each participant was required to have at least one time series from the language localizer task adapted from Pinel, et al. ^25^. This localizer was typically included alongside other functional tasks, depending on the specific project, whenever recording time allowed.
- fMRI time series data had to comprise at least 80 time points to be eligible for inclusion in the database.

After applying these selection criteria, the final dataset comprised 725 healthy participants with structural and functional MRI data (see Figure 1a). Participants’ sex and age were recorded at the time of registration. The neuroimaging analyses presented in the present paper, which include probabilistic atlas-based data, were conducted on the full sample (N = 725). Participants (389 women, 53.6%) ranged in age from 18 to 82 years (mean = 34; SD = 16.6; median = 27). Table 1 provides a demographic overview of the dataset. Additional demographic and behavioral measures are available for most participants, with a missingness rate that ranges between 1 and 52 %, depending on the variable (Table S1, S2). Moreover, 703 participants had further data on their language background (Figure 1b), and this subset was used to phenotype their linguistic profiles (reported in the section ***Phenotyping linguistic profiles***). Some individuals had multiple acquisitions (i.e., different StudyInstanceUID values), with 2 to 9 structural MRI (N = 157), and 2 to 14 fMRI runs each (N = 137), with a maximum of 6 runs per language. This subset was used exclusively to conduct data quality control through test-retest comparison. Last, for the analyses combining neuroimaging metrics with behavioral data (reported in the section ***Associations between brain structure and cognitive function accounting for sex and age***), we included a subset of 344 individuals (see Figure 1c) for whom there is a measure of cognitive function. The sample sizes indicated in brackets for each dataset refer to the subsets ultimately used in the presented analyses. Note that these figures may vary depending on the specific criteria applied during data filtering. To clarify the different subsets within NEUROLINGUA, an overview is provided in Figure 1.

**Figure 1.**
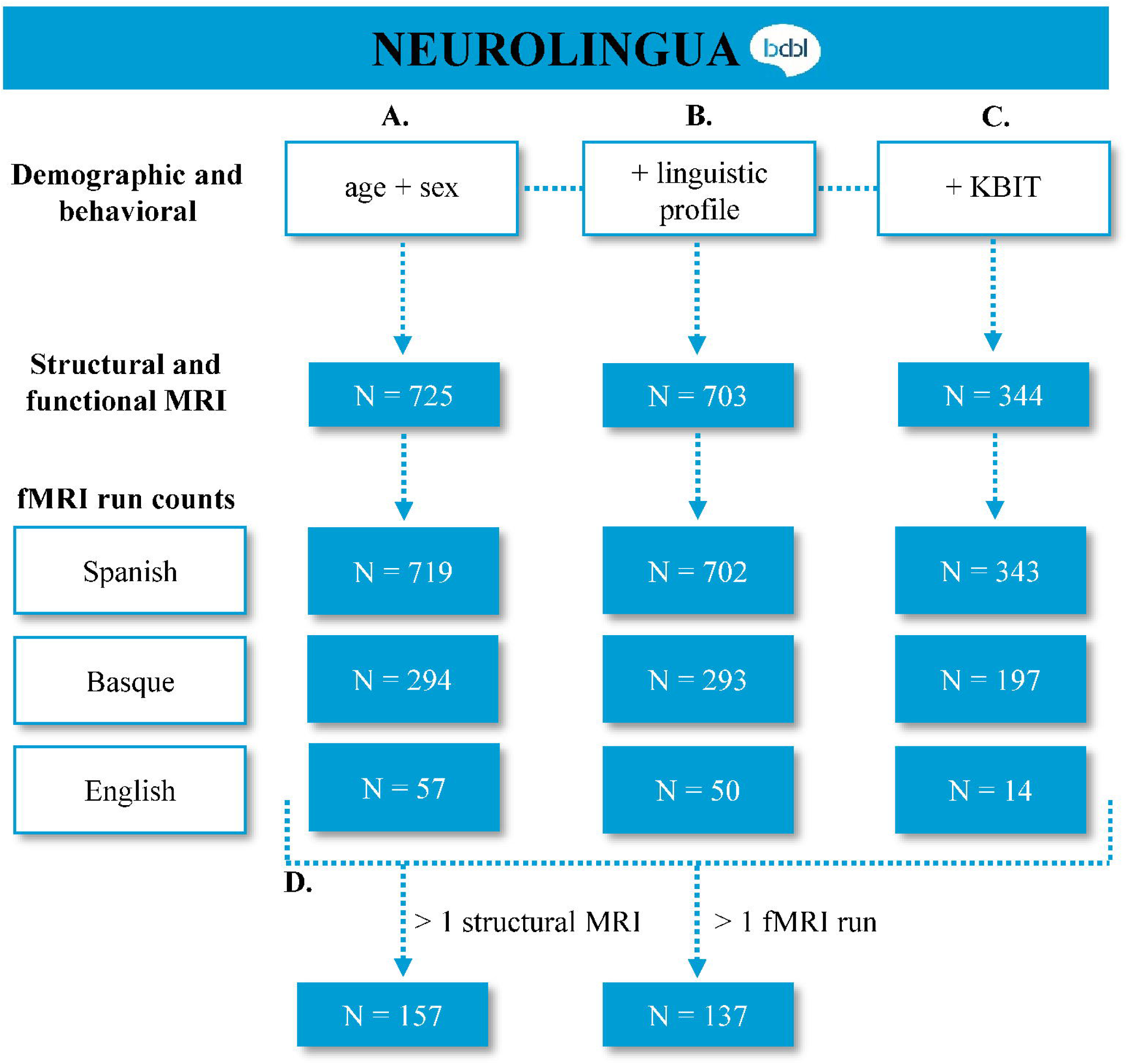
NEUROLINGUA dataset characterization. NEUROLINGUA database sample subsets (a, b, and c) and their corresponding sample sizes depending on the availability of additional data.

**Table 1.**
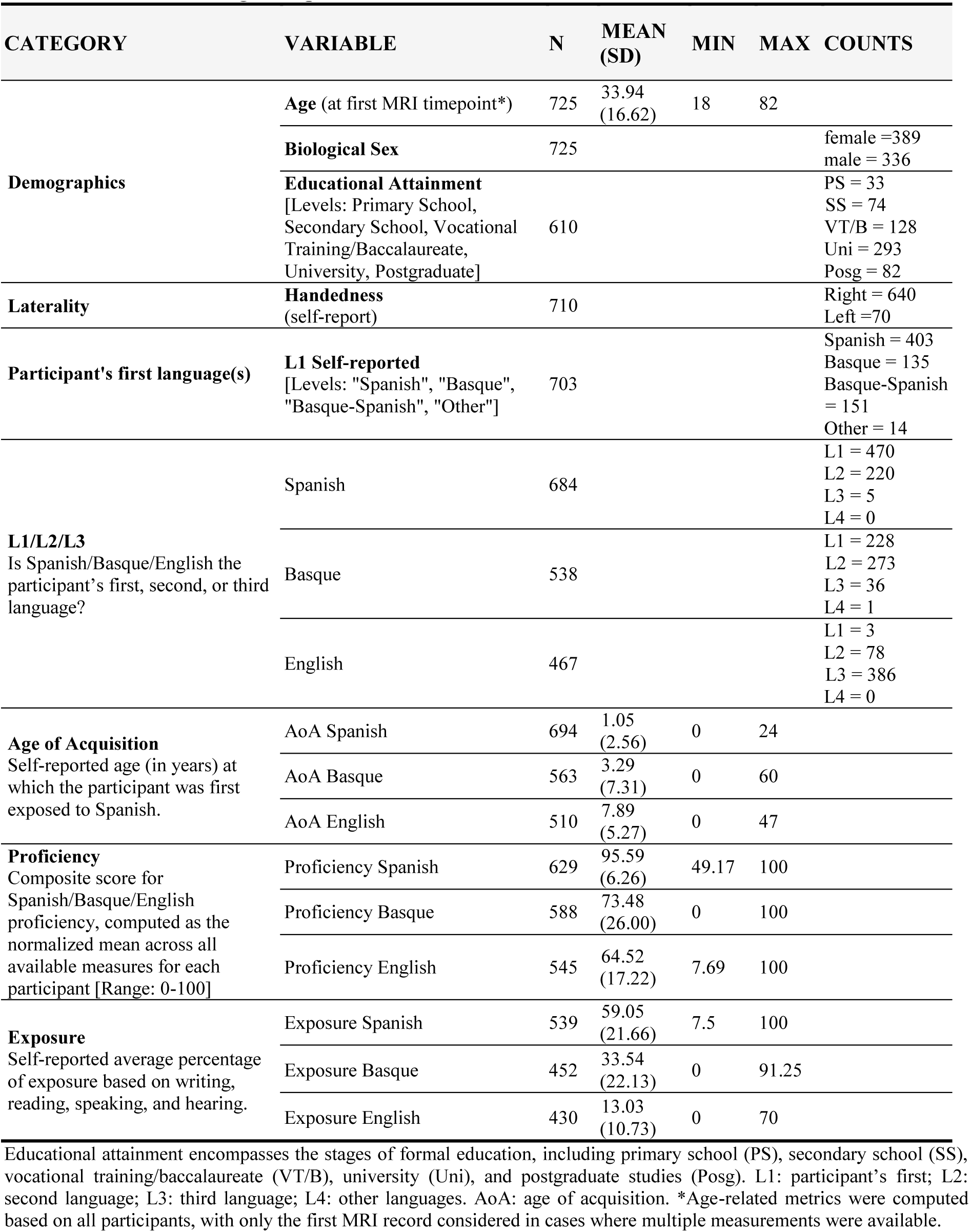
Descriptive statistics for demographic and linguistic variables (N_total_ = 725). The N reflects the non-missing data points for each variable.

**Table 2.** Cognitive assessment.

| DOMAIN | TASKS AND DESCRIPTION |
| --- | --- |
| **Verbal and non-verbal cognitive functioning** | **Intelligence test.** The Spanish version of the Kaufman Brief Intelligence Test (KBIT<sup>41</sup>) was used as an assessment of cognitive functioning. KBIT measures intelligence and includes 160 items divided into verbal and non-verbal knowledge. It has been linked to learning potential and cognitive reserve. The test provides three key values: verbal intelligence, non-verbal intelligence, and general intelligence. General intelligence is calculated as a composite measure derived from the scores of verbal and non-verbal intelligence. |
| **Language** | **Picture-naming test.** We used BEST<sup>29</sup>, a multilingual standardized battery for the assessment of language proficiency. It consisted of a picture-naming task composed of 65 (or 75) pictures. The naming task yields the percentage of correct responses (hits).<br>**Semi-structured interview.** A short interview guided by a multilingual linguist with experience in assessing language proficiency provided a score of the participants' language skills for each language, offering a more comprehensive index of linguistic competence, encompassing discourse-related aspects.<br>**Lexical decision task.** We adapted the Lextale<sup>30,42</sup>, a lexical decision task in which participants discriminated between words and pseudowords. The English version followed the original task design with 40 items, while the Basque and Spanish versions were expanded to 65 items each. For each language, we calculated the percentage of correct responses (hits). |

All individuals included in the database were residents of the Gipuzkoa province in the Basque Country at the time of data acquisition. Linguistically, this region is characterized by the use of both Basque and Spanish as primary languages. Additionally, English is widely spoken, particularly among younger adults. Consequently, the participants in this sample are immersed in a multilingual environment, ranging from near-monolingual individuals to those highly proficient in three or more languages. Our database provides a detailed linguistic characterization of the sample, making it possible to investigate the complex influence of bi(multi)lingualism on neural architecture.

### Demographic variables

The demographic variables collected include age, biological sex, and educational level. This information has been included in the files *behavioral_dataset.csv* and *participants.tsv*. In these files, age is expressed in years, and the computation of age did not take into account the day or month of birth; biological sex is expressed as a categorical variable that only considers the sex of the person at birth; and educational level is expressed as a categorical variable that reflects educational stages in Spain and can acquire the following values: “Primary”; “Secondary”; “Vocational Training/Baccalaureate”; “University”; and “Postgraduate”.

### Sociolinguistic and cognitive variables

The sociolinguistic variables consisted of measures of age of acquisition (AoA; self-reported), language exposure (%, self-reported), and four measures of language proficiency (self-reported level, interview-based level, picture naming score—BEST ^29^—, and lexical decision score—Lextale ^30^—), for three languages: Basque, Spanish, and English. The correlation matrix for these variables is shown in Figure S1.

Given the high correlation among the different proficiency measures within language (see Fig. S1), they were combined into a composite proficiency score per language, constructed by averaging all the available proficiency measures (in a common % scale) for each individual, thus capturing proficiency while minimizing missingness for this variable. Note that the lexical decision task was scaled so that chance level performance was coded as 0. To mitigate the extreme skewness of the AoA variables (long right-tail) and to ensure that all were in a common scale, we capped the maximum AoA values at 18 years (24 instances for self-reported AoA of Basque, 14 for English, and 1 for Spanish). As a result, we ended up with nine variables (AoA, exposure, and proficiency for each of the three languages) that capture most of the variability in linguistic profiles. These variables were then combined to create an index for bilingualism and another for trilingualism, as described below and defined in Table 3.

**Table 3.** Overview of linguistic variables. Bilingualism and Trilingualism indices were created based on AoA, proficiency, and exposure variables in Spanish, Basque, and English.

| MEASURES<br>unit [range] |  | AGE OF<br>ACQUISITION (AoA)<br>Years [0-18] | LANGUAGE<br>PROFICIENCY<br>Composite score [0,100] | LANGUAGE<br>EXPOSURE<br>% [0,100] |
| --- | --- | --- | --- | --- |
| Languages | Spanish | N = 629 | N = 627 | N = 505 |
|  | Basque | N = 522 | N = 587 | N = 417 |
|  | English | N = 509 | N = 543 | N = 430 |
| Scaling factor |  | 18 | 100 | 100 |
| Distance | *distance* | *Basque – Spanish* | *Spanish – Basque* | *Spanish – Basque* |
|  | *scaled* | 18 | 100 | 100 |
| Bilingualism Index<br>[range -1; 1] | | | $\frac{1}{var} \sum_{var=1}^m distance_{var}$ | |
| Scaling Factor | | 18 | 100 | $\frac{100}{n}$ |
| Mean per measure across<br>available languages (n) | | $\frac{\frac{1}{n} \sum_{i=1}^n 18 - AoA_i}{ScalingFactor}$ | $\frac{\frac{1}{n} \sum_{i=1}^n Proficiency_i}{ScalingFactor}$ | $1 - \frac{\frac{1}{n} \sum_{i=1}^n Exposure_i}{\frac{100}{ScalingFactor}}$ |
| Trilingualism Index [range:<br>0-3] | | | $n \times \frac{1}{var} \sum_{var=1}^m mean_{var}$ | |

n = number of available languages for a given individual; m = number of available variables for each individual.

#### Spanish-Basque Bilingualism Index

To capture variability in linguistic profiles for Spanish and Basque (the primary languages among participants), we developed a Spanish-Basque Bilingualism Index. First, we calculated difference scores between the two languages for Age of Acquisition (AoA, Basque-Spanish), exposure (Spanish-Basque), and proficiency (Spanish-Basque). Each score was then scaled by dividing by its maximum possible value (100 for exposure and proficiency, 18 for AoA). We averaged all available standardized differences for each individual, resulting in an index ranging from −1 to 1 for participants with measurements in both Spanish and Basque for at least one variable. To maximize the sample, if an individual did not have one of the variables, the index was computed by averaging the rest of the available variables. An index of −1 indicates a Basque-dominant profile, 0 a balanced Basque-Spanish profile, and 1 a Spanish-dominant profile, reflecting each participant’s balance or dominance in Basque or Spanish (see Table 3).

#### Spanish-Basque-English Trilingualism Index

To assess trilingualism in this language combination, we constructed a second index that includes measures for Spanish, Basque, and English. (Other language combinations were not considered due to insufficient data.) We began by calculating an average score across languages for each measure:

- Proficiency: Composite score computed as the scaled mean proficiency across all possible available information per participant (interview, lexical decision task, and picture naming task, among others).
- AoA: The mean distance to 18 (i.e., maximum possible AoA), calculated as mean(18 - AoA.lang), normalized by 18.
- Exposure: Given that exposure is recorded as a percentage, we defined balanced exposure relative to the number of languages (n). For instance, balanced exposure for two languages is 50%, and for three languages, 33.33%. We then calculated each person’s mean absolute deviation from this balanced exposure across languages and scaled it by 100/n to achieve values from 0—closest to balanced exposure— to 1—furthest from balanced exposure. Finally, we inverted this score (1 - index) so that 0 represents a monolingual experience and 1 represents a balanced multilingual experience.

Each of these measures yielded an index between 0 and 1, where 0 denotes a monolingual profile, and 1 represents a balanced, proficient experience across multiple languages. The three indices (proficiency, AoA, exposure) were then averaged, and the mean was multiplied by n (1 to 3) to adjust for each participant’s number of languages. The resulting Trilingualism Index ranges from 0 to 3, with each point indicating a balanced, complete experience with an additional language (1 = monolingual, 2 = balanced bilingual, 3 = balanced trilingual). These values are general indicators, as an index of 2 may reflect full proficiency in one language and intermediate proficiency in two others; scores below 1 indicate incomplete proficiency/exposure in one (or possibly more) language(s).

### Description of the functional localizer

For this study, we used a version of the functional language localizer originally designed by Pinel, et al. ^25^ adapted for Spanish, Basque, and English. This localizer was developed to identify, at the individual level, the functional network that supports language comprehension. As in the original version, the timing and distribution of experimental trials were optimized to maximize the contrast between different experimental conditions by using a standard optimization genetic algorithm ^31^. The task included a mixture of visual and auditory trials, presented randomly in equal distribution. The trials followed a nested design, with auditory and visual sentences categorized into three experimental conditions based on the task to be performed: (1) sentences for passive comprehension; (2) sentences involving arithmetic problem-solving; and (3) sentences requiring motor action response. Additionally, the task included a control condition involving the presentation of flashing horizontal or vertical checkerboards. To ensure correct task performance, participants received written instructions before the task began. Each trial started with a jittered fixation point, lasting between 200 ms and 6600 ms. To optimize visual stimulus presentation, each visual sentence was divided into four segments of three to four words each. These segments were shown successively for 250 ms each, with 100 ms intervals between them, resulting in a total presentation time of 1.5 seconds per trial. The auditory trials, recorded by a male speaker, had a mean duration of 2.45 seconds. Figure 2 illustrates the experimental design, including examples of the trials for each experimental condition.

**Figure 2.**
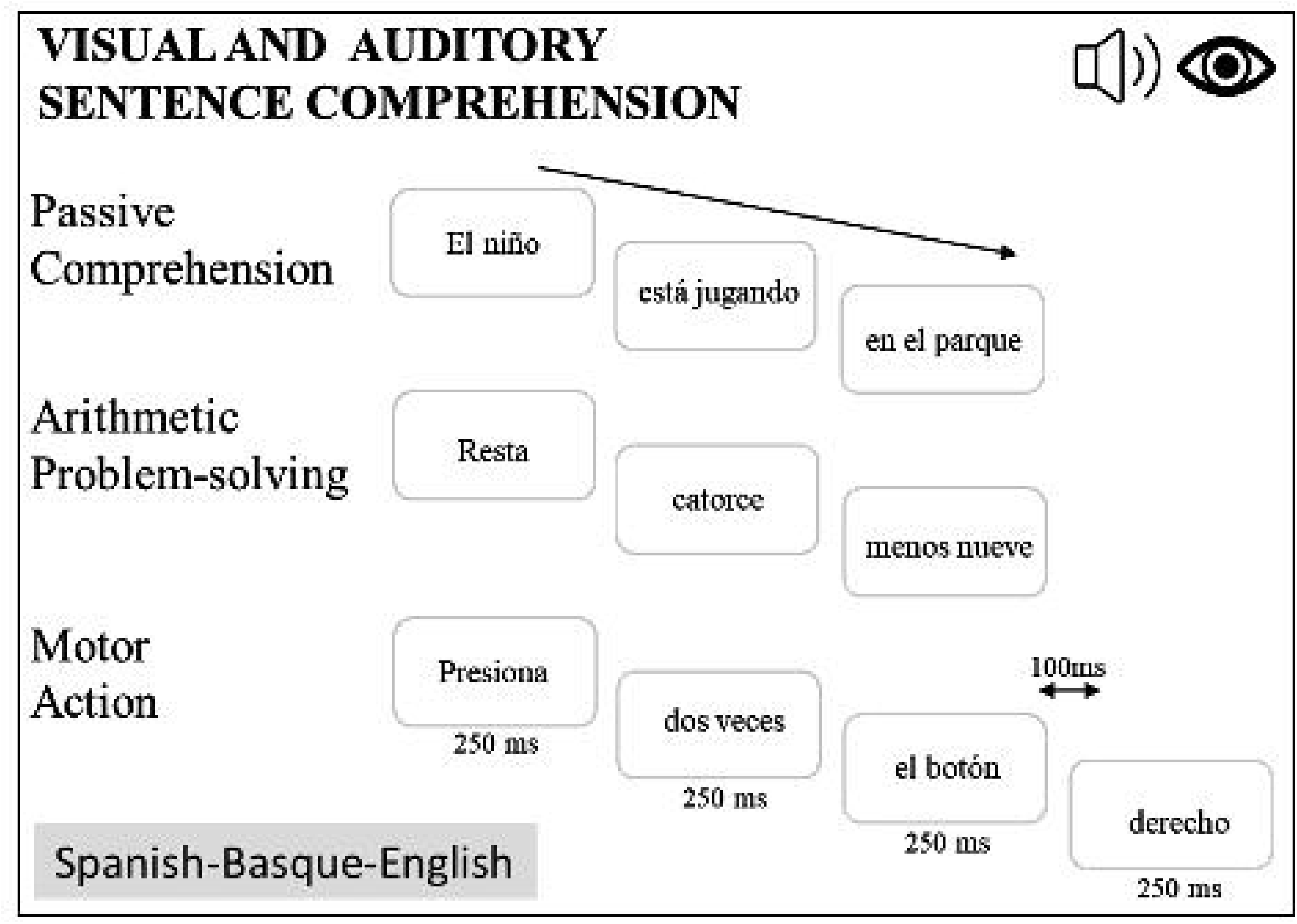
Task-based fMRI Design. The trials followed a nested design, with auditory and visual sentences categorized into three experimental conditions based on the task performed: passive comprehension (e.g., *“El niño está jugando en el parque”* [The boy is playing in the park]), arithmetic problem-solving (e.g., *“Resta catorce menos nueve”* [Subtract nine from fourteen]), and motor action execution (*“Presiona dos veces el botón derecho”* [Press two times the right button]). Adapted from the functional language localizer designed by Pinel, et al. ^25^.

### f/MRI data acquisition

All participants underwent an MRI session in a 3T Siemens Magnetom scanner (Siemens AG, Erlangen, Germany). High-resolution T1-weighted images were acquired with a 3D MPRAGE sequence using a 64-channel head coil. The imaging protocol primarily used a Repetition Time (TR) of 2530 ms, a flip angle of 7°, and a voxel resolution of 1×1×1 mm³, resulting in a total of 785 images. Of these, 481 images were acquired with an Echo Time (TE) of 2.36 ms, utilizing 176 contiguous sagittal slices, 256 coronal slices, and 256 axial slices. The remaining 462 images were obtained with a TE of 2.97 ms, comprising 256 contiguous sagittal slices, 256 coronal slices, and 176 axial slices. Additionally, several other parameter configurations were used for image acquisition, with detailed information on these settings and the corresponding image counts provided in Table 4. The structural images for three participants (sub-0193, sub-0222, sub-0259) were used solely for normalizing the functional data to MNI space. These T1-weighted scans were acquired with parameters that do not meet the quality standards required for morphometric feature extraction. As a result, these participants were excluded from subsequent structural and structure-function relationship analyses.

**Table 4.** Parameters of T1-weighted MRI sequences. These parameters were optimized to ensure high-resolution anatomical imaging suitable for subsequent neuroimaging analyses.

| **ECHO TIME**<br>**(ms)** | **REPETITION**<br>**TIME (ms)** | **FLIP ANGLE**<br>**(grades)** | **BASE**<br>**RESOLUTION** | **SLICE**<br>**THICKNESS**<br>**(mm)** | **COUNT**<br>**(n)** |
| --- | --- | --- | --- | --- | --- |
| 2.36 | 2530 | 7 | 256 | 1 | 481 |
| 2.97 | 2530 | 7 | 256 | 1 | 462 |
| 2.97 | 2300 | 9 | 256 | 1 | 17 |
| 3.50 | 2530 | 7 | 256 | 1 | 15 |
| 2.34 | 2400 | 8 | 320 | 0.7 | 43 |

Functional images were primarily acquired using a T2*-weighted gradient-echo echo-planar imaging sequence with two main sequence configurations. The first parameter set used for 733 acquisitions involved 406 scans with a TE of 35 ms, a TR of 850 ms, a flip angle of 56°, and a FOV of 212 x 212 mm^2^. This configuration covered 66 slices, each with a voxel size of 2.4 × 2.4 × 2.4 mm³. The effective echo spacing was 0.50 ms, with an acceleration factor of 6, and a matrix size of 88 × 88. The second most commonly used parameter set used for 674 datasets comprised 145 scans with 39 slices and a voxel size of 3 × 3 × 3 mm³, with a TE of 30 ms, a TR of 2400 ms, an acceleration factor of 6, and a flip angle of 90°. This configuration also had an effective echo spacing of 0.51 ms, and a FoV of 192 x 192 mm^2^, with a matrix size of 64 × 64. While these two configurations were the most frequently used, a variety of additional imaging parameters were also employed for the acquisition of fMRI data. Details on these configurations and their respective parameters are provided in Table 5.

**Table 5.**
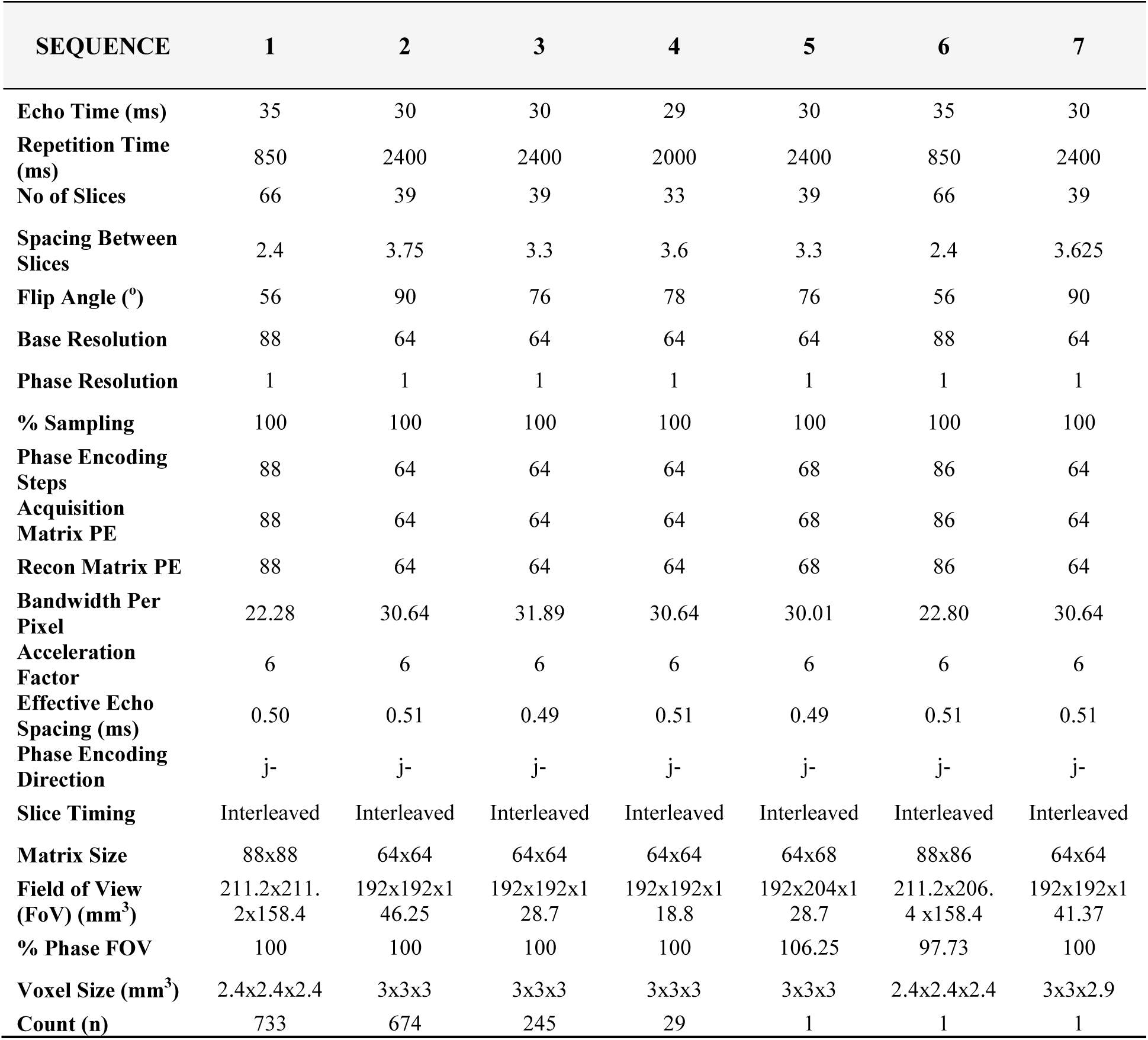
Parameters of T2* weighted functional MRI sequences.

| SEQUENCE | 1 | 2 | 3 | 4 | 5 | 6 | 7 |
| --- | --- | --- | --- | --- | --- | --- | --- |
| Echo Time (ms) | 35 | 30 | 30 | 29 | 30 | 35 | 30 |
| Repetition Time (ms) | 850 | 2400 | 2400 | 2000 | 2400 | 850 | 2400 |
| No of Slices | 66 | 39 | 39 | 33 | 39 | 66 | 39 |
| Spacing Between Slices | 2.4 | 3.75 | 3.3 | 3.6 | 3.3 | 2.4 | 3.625 |
| Flip Angle (°) | 56 | 90 | 76 | 78 | 76 | 56 | 90 |
| Base Resolution | 88 | 64 | 64 | 64 | 64 | 88 | 64 |
| Phase Resolution | 1 | 1 | 1 | 1 | 1 | 1 | 1 |
| % Sampling | 100 | 100 | 100 | 100 | 100 | 100 | 100 |
| Phase Encoding Steps | 88 | 64 | 64 | 64 | 68 | 86 | 64 |
| Acquisition Matrix PE | 88 | 64 | 64 | 64 | 68 | 86 | 64 |
| Recon Matrix PE | 88 | 64 | 64 | 64 | 68 | 86 | 64 |
| Bandwidth Per Pixel | 22.28 | 30.64 | 31.89 | 30.64 | 30.01 | 22.80 | 30.64 |
| Acceleration Factor | 6 | 6 | 6 | 6 | 6 | 6 | 6 |
| Effective Echo Spacing (ms) | 0.50 | 0.51 | 0.49 | 0.51 | 0.49 | 0.51 | 0.51 |
| Phase Encoding Direction | j- | j- | j- | j- | j- | j- | j- |
| Slice Timing | Interleaved | Interleaved | Interleaved | Interleaved | Interleaved | Interleaved | Interleaved |
| Matrix Size | 88x88 | 64x64 | 64x64 | 64x64 | 64x68 | 88x86 | 64x64 |
| Field of View (FoV) (mm<sup>3</sup>) | 211.2x211.2x158.4 | 192x192x146.25 | 192x192x128.7 | 192x192x118.8 | 192x204x128.7 | 211.2x206.4x158.4 | 192x192x141.37 |
| % Phase FOV | 100 | 100 | 100 | 100 | 106.25 | 97.73 | 100 |
| Voxel Size (mm<sup>3</sup>) | 2.4x2.4x2.4 | 3x3x3 | 3x3x3 | 3x3x3 | 3x3x3 | 2.4x2.4x2.4 | 3x3x2.9 |
| Count (n) | 733 | 674 | 245 | 29 | 1 | 1 | 1 |

### Micro- and macrostructural MRI extraction

To ensure an efficient and streamlined analysis workflow, the CAT12 toolbox ^1^ was utilized for estimating micro- and macrostructural properties. This toolbox is an extension of SPM12 (https://www.fil.ion.ucl.ac.uk/spm/software/spm12/) written in MATLAB. Specifically, voxel-based morphometry (VBM) was used, a semi-automated neuroimaging technique widely employed to quantify macrostructural changes in brain volume and density. A standardized in-house processing pipeline was applied to the T1-weighted MRI images. The pipeline begins with the application of a spatially adaptive nonlocal means (SANLM) denoising filter ^32^ to enhance image quality, followed by internal resampling to manage low-resolution images and anisotropic spatial resolutions. Next, data bias correction and affine registration are performed to optimize the outcomes of subsequent processing steps, followed by unified SPM segmentation ^33^. This step provides initial estimates that guide the subsequent refined voxel-based processing.

Refined voxel-based processing continues with skull stripping and the division of the brain into left and right hemispheres, subcortical regions, and the cerebellum. The detection of local white matter hyperintensities was also performed at this stage and was accounted for during spatial normalization. A local intensity transformation of all tissue classes follows, which helps mitigate the impact of higher gray matter intensities. The final segmentation step involves a maximum adaptive a posteriori segmentation approach, refined with partial volume estimation to accurately determine the fractional content of each tissue type per voxel. Subsequently, tissue segments—including gray matter (GM), white matter (WM), and cerebrospinal fluid (CSF)—are spatially normalized to a common reference space using Geodesic Shooting techniques ^34^. During this process, brain volume metrics such as Total Intracranial Volume (TIV), GM, WM, and CSF were calculated from the T1-weighted images.

In parallel, microstructural properties were extracted following the recommendations provided by https://neuro-jena.github.io/cat12-help/. Roughly, surfaces were created using a projection-based thickness method, which estimates the initial cortical thickness and the initial central surface. After this initial step, topological defects are corrected, resulting in final refined surfaces corresponding to the central, pial, and white surfaces. Subsequently, the pial and white surfaces are used to refine the initial cortical thickness estimate using the FreeSurfer thickness metric ^35^. The final central surface was used to calculate cortical folding metrics. Finally, a surface registration step was performed, where individual central surfaces are spherically inflated following a minimal distortion algorithm. This step results in a one-to-one mapping between the folding patterns of the individual spheres and the template using a two-dimensional (2D) version of the DARTEL approach ^34^.

### Functional MRI preprocessing and statistical analysis

Functional event-related data were analyzed using SPM12 and related toolboxes (i.e., RobustWLS - Robust Regression using Weighted-least-squares: http://www.diedrichsenlab.org/imaging/robustWLS.html and MarsBar - MARSeille Boite A Region d’interet: http://marsbar.sourceforge.net/). Raw functional scans were slice-time corrected, taking the middle slice as reference, spatially realigned, unwarped, and coregistered with the anatomical T1 and normalized to MNI space using the unified normalization segmentation procedure ^33^. Global effects were then removed using a global signal regression analysis ^36^, after which the data were smoothed using an isotropic Gaussian kernel with a full-width half maximum of 8 mm. Statistical parametric maps were generated using a univariate general linear model with a regressor for each stimulus type obtained by convolving the canonical hemodynamic response function with delta functions at stimulus onsets, also including the six motion-correction parameters as regressors of no interest. The stimulus onsets included seven different components. The first corresponded to the onset of each sentence trial and was modeled as a single regressor, independently of the experimental conditions. The next six corresponded to the three experimental conditions (Passive Comprehension, Arithmetic Problem Solving, and Motor Action), accounting for the modality (visual and auditory). GLM parameters were estimated using a robust regression approach with weighted least squares ^37^.

Following the methodology proposed by Lipkin, et al. ^26^ and utilizing an in-house custom code available at OSF, we computed the overlap of individual activation maps for the contrasts auditory comprehension > baseline and visual comprehension > baseline across all participants. Specifically, we used whole-brain t-maps generated from the first-level analysis.

For each individual t-map, we identified the top 10% of voxels with the highest t-values. As in Lipkin, et al. ^26^, we then binarized the maps using this threshold (90th percentile). These binary maps were summed and divided by the total number of participants, yielding an atlas in which each voxel had a value between 0 and 1. This value represents the probability of voxel activation relative to the baseline, with 1 indicating activation in 100% of participants and 0 indicating no activation in any participant. This approach allowed us to assess the consistency of our findings with those of Lipkin, et al. ^26^ (see Figures 5 and 6). However, since the 90th percentile of the t-map distribution may result in low thresholds that do not correspond to significantly activated voxels, we implemented an alternative method based on false discovery rate (FDR) correction, following Benjamini and Hochberg’s procedure. Instead of using the 90th percentile as a threshold, we computed p-values from the same t-maps and combined them using Stouffer’s method. We then applied FDR correction to determine the threshold. Using this approach, we generated an alternative atlas with the same characteristics as in Lipkin, et al. ^26^, ensuring a more statistically robust selection of activated voxels (Figures S5 and S6).

### f/MRI visualization App: NEUROLINGUA sMRI vizApp

The MRI structural and functional derivatives have also been included in an interactive shiny app created with R’s “shiny” package (R version 4.1.3, *shiny* v. 1.7.2, *plotly* v4.10.4), which can be explored for quality control and familiarization with the data (https://www.bcbl.eu/databases/shiny/neurolinguapp). This application displays descriptive statistics of the structural and functional MRI data for 16 different atlases extracted with CAT12, allowing the exploration of specific regions by checking their distribution and relationship with age. Moreover, it also provides additional analyses for the subjects that have multiple sessions, by showing correlations across sessions, and test-retest correlations between specified sessions.

## Data Record

MRI/fMRI data in DICOM format were converted to the Brain Imaging Data Structure (BIDS) standard using BIDScoin (https://doi.org/10.3389/fninf.2021.770608) ^38^. Per BIDS’ hierarchical structure, each participant’s data is stored in a subfolder labeled with a numerical code generated through a double-blind process. Neuroimaging data were stored as compressed NIfTI files (*.nii.gz), with acquisition parameters for each scan saved in accompanying JSON files (.json), excluding any personally identifiable details. Within each participant’s folder, two subfolders are created: *“anat/”* and *“func/”*, originally containing the raw structural and functional images, respectively. Specifically, the *“anat/”* folder contains defaced structural images derived from the *pydeface* function implemented in BIDS and the transformation matrices necessary for transitioning between native and normalized MNI spaces. The *“func/”* folder holds functionally preprocessed images, with smoothing as the final preprocessing step. To enhance reproducibility and transparency, the dataset derived from the analyses is publicly accessible via the OpenNeuro platform (https://openneuro.org/). Figure 3 illustrates the folder structure on the platform. This approach ensures compliance with personal data protection laws by safeguarding participants’ identities while also promoting the reproducibility of the exploratory analyses presented in this study. Event details for functional image time series are provided in the *_events.tsv file. Sociolinguistic, sociodemographic, and cognitive data, along with indices derived from these variables, are included in the behavioral_dataset.csv file in the main directory.

**Figure 3.**
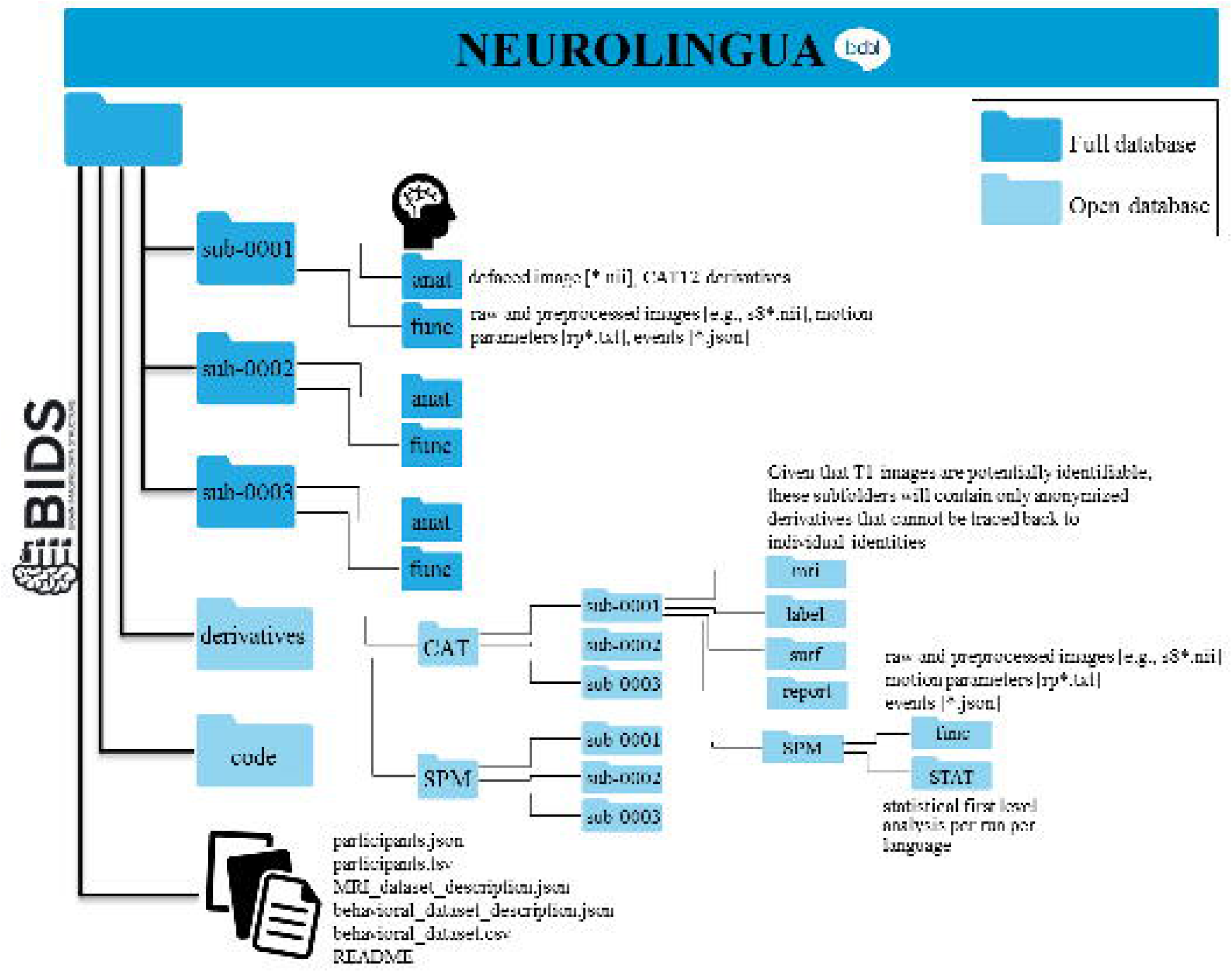
Database folder organization. Folders in dark blue will be shared upon request, while those in light blue are openly accessible via OpenNeuro.

Identifiable T1-weighted raw data will be made available upon request (https://www.bcbl.eu/DataSharing/Neurolingua/) since consent for publicly sharing potentially identifiable data was not obtained at the time of data collection. Access will require signing a confidentiality agreement between the applicant and BCBL, specifying the terms of data use. All other neuroimaging data (including raw functional data and MRI/fMRI derivatives) is openly available on the OpenNeuro platform in anonymized form. All personally identifiable information from behavioral and neuroimaging data was removed. This included purging the date, time, participant’s weight, height, and day and month of birth. Basic demographic information, such as age and biological sex, is included in *participants.tsv* file in the main BIDS directory.

## Technical Validation

While we have taken great care to produce a fully consistent dataset, a small margin of error may still be present. These errors stem from various factors, including technical malfunctions, participant non-compliance, and occasional human error. However, we are confident that these minor issues do not compromise the overall value of the data provided in NEUROLINGUA. The exploratory analyses presented below aim to illustrate the potential applications of the database, as well as possible sources of bias.

### Sample composition

Demographic, sociolinguistic, and cognitive data were available for all participants, though the extent of available data varied based on the specific projects in which they participated. To establish a consistent linguistic profile across all participants, we harmonized these variables, resulting in a set of 48 variables used for analysis. The selected variables, categorized as shown in brackets, spanned eight categories: Demographics (4), AoA (3), Language Exposure (11), L1/L2/L3 (4), Language Proficiency (15), Cognition (4), Laterality (3), and MRI Data (4). Table S3 presents the linguistic profiles of the participants. Please note that the number of variables may differ per participant, as not all variables were mandatory or applicable (e.g., whether a participant had acquired a third language). Key demographic details for the dataset are illustrated in Figure 4.

**Figure 4.**
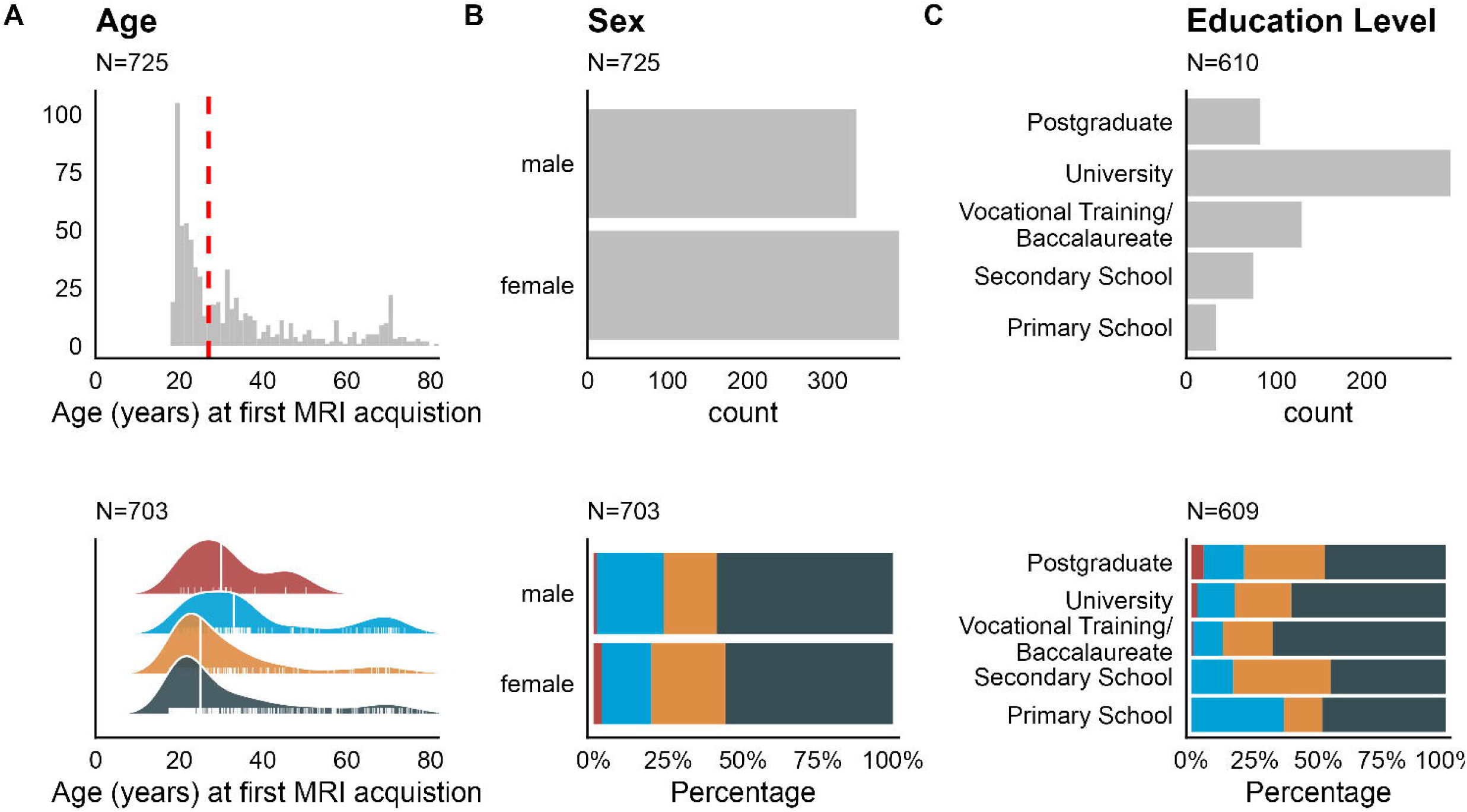
Demographic variables. The graph includes the distribution by (A) age, (B) sex, and (C) education level. The upper panel shows the number of participants for each variable. The lower panel the distribution of each variable per L1. The dashed red line (upper panel) and solid white lines (lower panel) indicate the median value for each distribution. Age-related metrics were computed based on all participants, with only the first MRI record considered in cases where multiple measurements were available.

### Phenotyping linguistic profiles

The analysis of estimated bilingualism and trilingualism indices reveals that NEUROLINGUA encompasses a broad range of linguistic profiles, from near-monolingual Spanish speakers to bilingual individuals with L1 dominance, and extending to non-dominant bilingual speakers. Figures 5A and 5B present the histograms for each profile and index, while Figure 5C illustrates that, despite some overlap, the indices effectively capture distinct boundaries between these profiles, as confirmed by a k-means clustering (R package *factoextra*, v. 1.0.7) analysis that identified four clusters (Figure 5E, between_SS / total_SS = 78.7%, average Silhouette width = 0.45) mapping roughly into the L1 categories (χ^2^(9) = 370.83, p-value = 2.28×10^-74;^ Figure 5D). Cluster 1 (N = 243) consisted of the most multilingual and Spanish-Basque balanced people (L1: Basque/Spanish = 44%, Spanish = 34%, Basque = 21%), cluster 2 (N = 68) consisted of most Basque-dominant people (L1: Basque = 90%, Basque/Spanish = 6%, Spanish = 4%), cluster 3 (N = 110) was most Spanish-dominant and less multilingual (L1: Spanish = 95%, Other = 4%, Basque/Spanish = 2%) and cluster 4 (N = 202) were Spanish-dominant but more multilingual (L1: Spanish = 70%, Basque/Spanish = 18%, Basque = 8%, Other = 3%).

**Figure 5.**
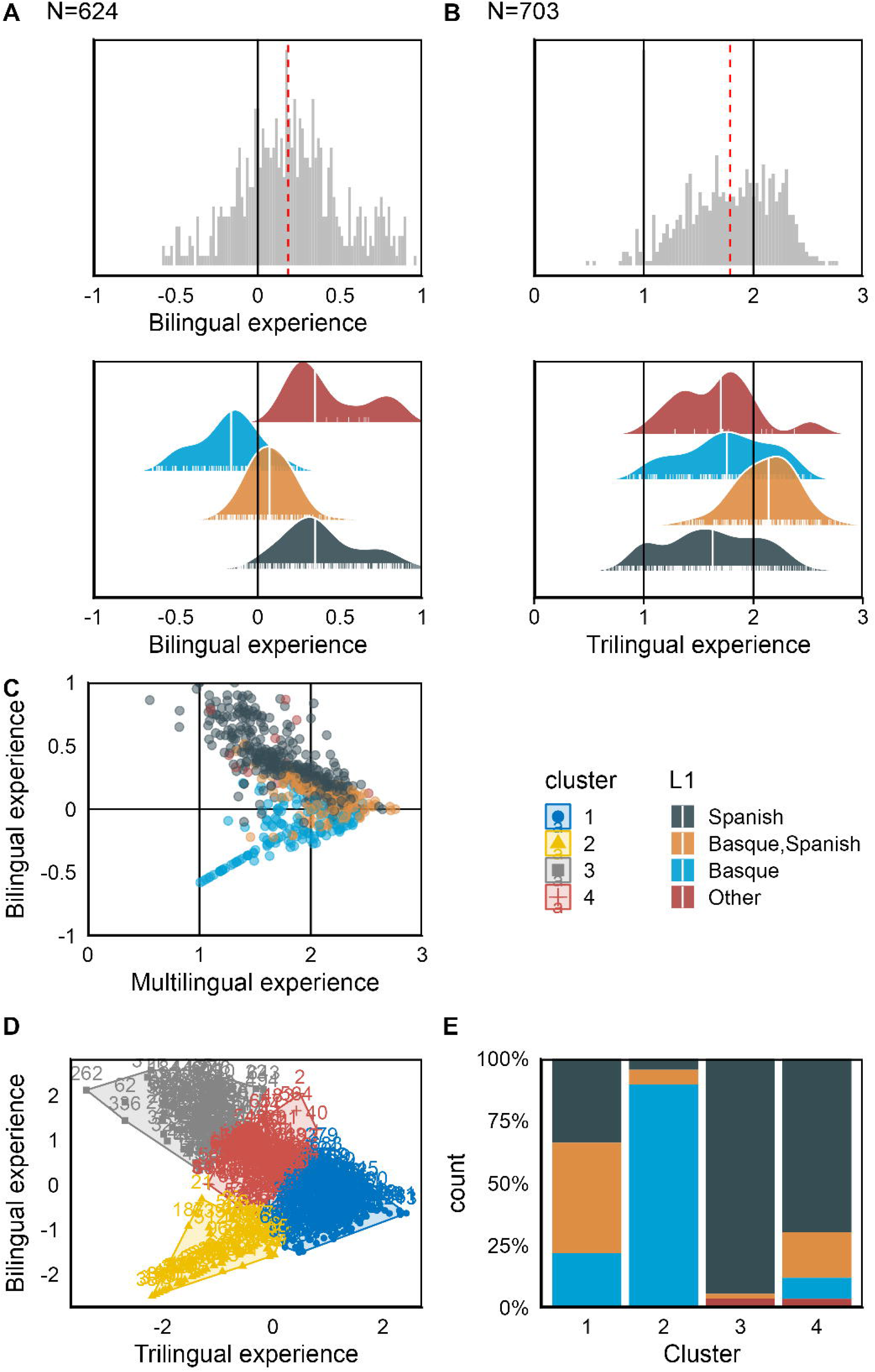
Phenotyping linguistic profiles. Distributions of the linguistic indices: A) Bilingualism index and B) Trilingualism index. C.) Relationship between the indices grouped by L1. D) Cluster assignment per L1. E) Cluster plot.

### Structural MRI validation

Microstructural and macrostructural properties were estimated using all images in the database, encompassing cases with multiple acquisitions per participant. Alongside volumetric images and refined surfaces, we provided *sMRI*.csv tables containing estimated values for each region of interest (ROI), as defined by various atlases. The metrics used in the analyses presented in this paper are specifically derived from the AAL3 atlas ^39^. These tables also include quality metrics, such as the ratio of the weighted overall image quality (IQR) to the mean quartic Z-score, indicating signal-to-noise ratio. To validate our structural measurements, we compared cortical thickness and total volume values against those provided by Bethlehem, et al. ^40^ in their interactive open resource for cross-database morphometric comparisons. This comparison positions our results within the mean of the distribution reported in the tool (Figures S2, S3).

### Functional MRI validation

Analysis of the fMRI data following the approach proposed by ^26^ confirmed NEUROLINGUA’s effectiveness in localizing the language comprehension network. Conjunction analyses allowed us to characterize how information flow within the language network varies by modality. As shown in Figures 6 and 7, the task successfully alternated between primary and secondary visual regions for visual conditions and primary and secondary auditory regions for auditory conditions. For results of the conjunction analysis using the alternative methodology described in the section ***Functional MRI preprocessing and statistical analysis*** where individual-level thresholds were determined using FDR correction, refer to supplementary figures S4 and S5. Figure S6 illustrates individual variability in the general contrasts, showing that the regions with the highest T values (mean T value per run) contrasts correspond to the expected modality-dependent regions, occipital regions for “Visual” (all Visual runs vs baseline), and temporal regions for “Auditory” (all auditory runs vs baseline). Figure 8 displays the mean of the individual maximum T-values within each of the identified language hubs and their contralateral counterparts. As shown, all regions previously defined as part of the language network exhibit significant activation compared to the null hypothesis (µ = 0) in both the visual and auditory contrasts. While the distribution of activation may vary depending on modality, all key areas—including the inferior frontal gyrus (pars orbitalis, triangularis, and opercularis), middle and superior temporal regions, supramarginal, angular, and inferior parietal gyri— demonstrate significant and individually replicable recruitment.

**Figure 6.**
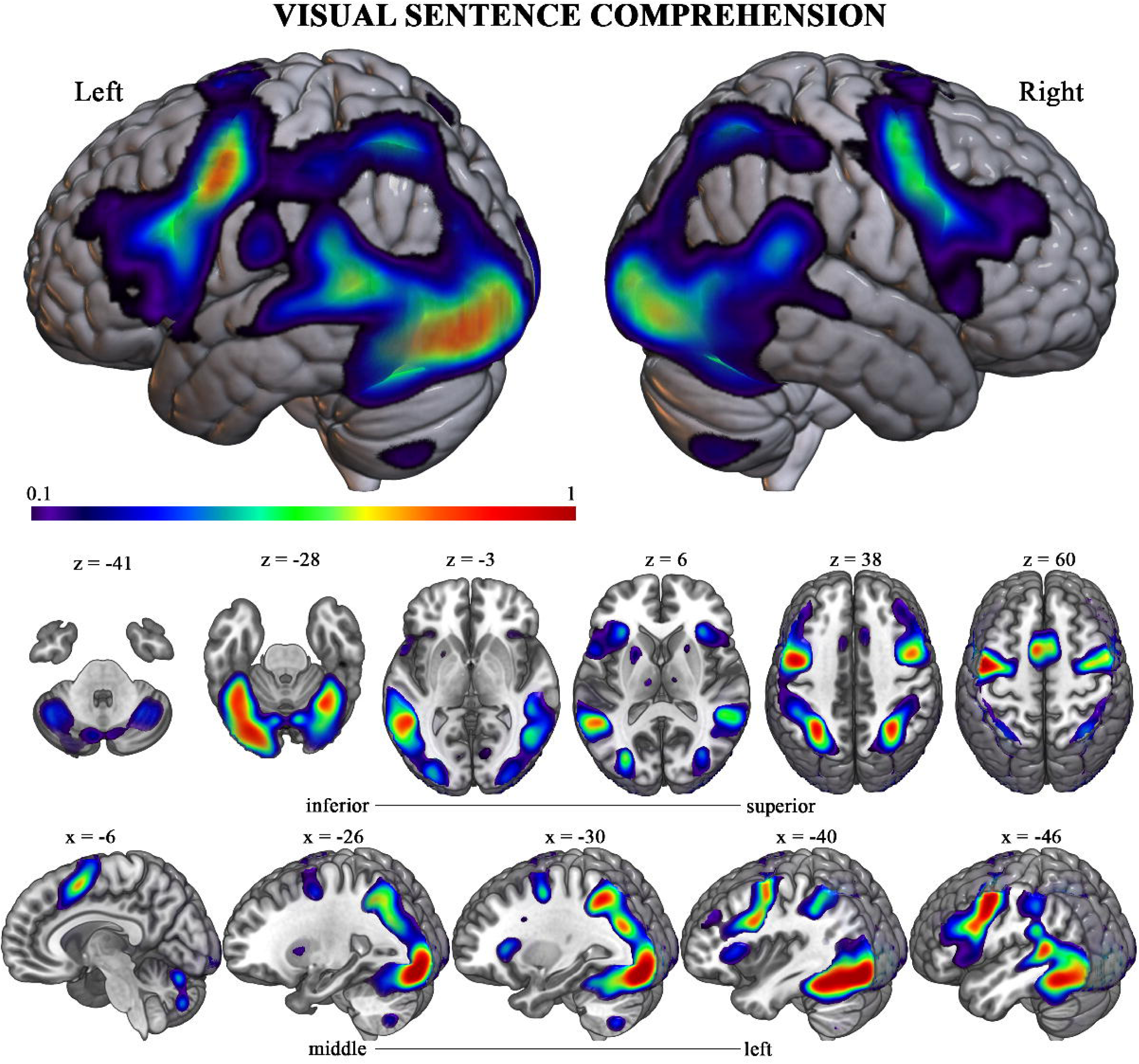
Visual language comprehension functional topography. Brain renderings and sagittal and axial sections showing the probabilistic functional activation for the “visual language comprehension” > baseline contrast based on overlaid individual binarized activation maps (where in each map, the top 10% of voxels are selected, as described in the text).

**Figure 7.**
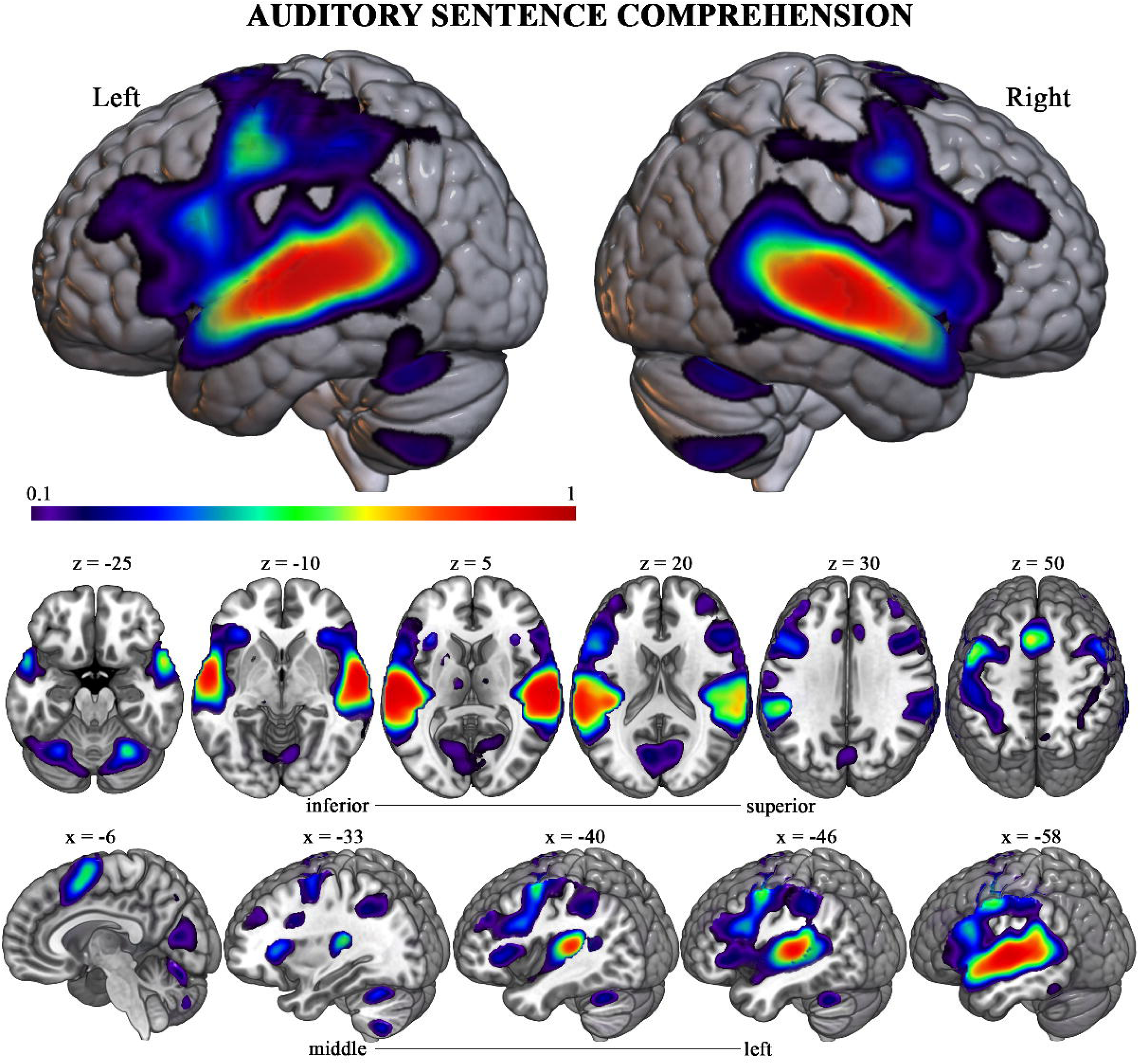
Auditory language comprehension functional topography. Brain renderings and sagittal and axial sections showing the probabilistic functional activation for the “auditory language comprehension” > baseline contrast based on overlaid individual binarized activation maps (where in each map, the top 10% of voxels are selected, as described in the text).

**Figure 8.**
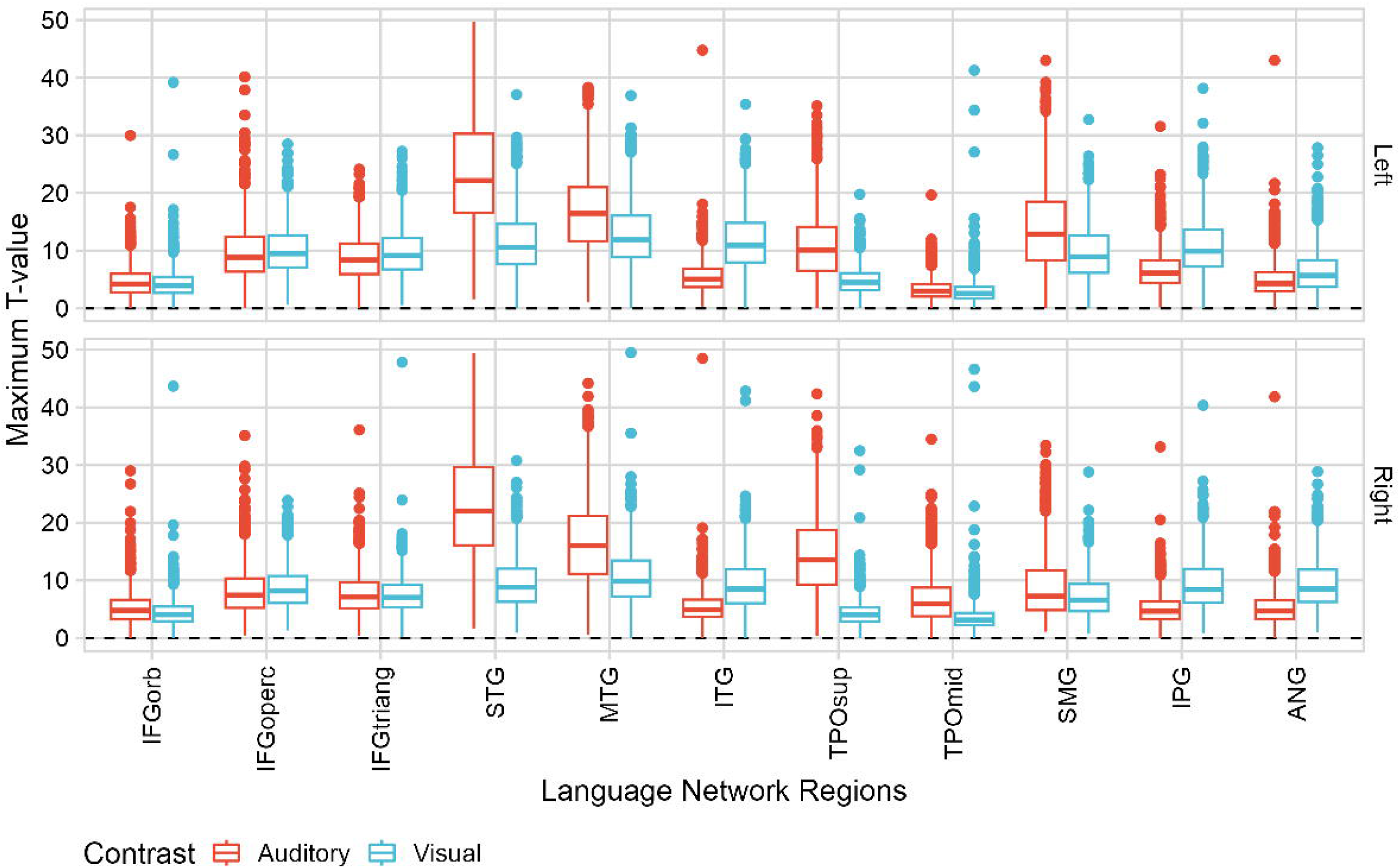
Maximum T-values for auditory and visual contrasts (compared to the null hypothesis) across predefined language hubs. Language hubs were defined based on the literature, with regions of interest derived from the AAL3 atlas. Each boxplot shows the mean and standard deviation of activation per region, across individuals and language runs. The upper panel displays regions in the left hemisphere; the lower panel shows their contralateral homologues. Note that mean activation in each region exceeds zero, indicating significant recruitment across conditions. Note that maximum T-values above 50 have been excluded for visualization purposes.

### Associations between brain structure and cognitive function, accounting for sex and age

This analysis included participants who completed the Kaufman Brief Intelligence Test, (KBIT ^1,2^), used to estimate verbal and nonverbal cognitive abilities. The KBIT is commonly employed as a screening tool to provide a general estimate of intellectual functioning. Specifically, we conducted multiple regression analyses with age, sex, and cognitive function (i.e., KBIT-derived scores) as additive predictors of global structural brain measures, including total gray matter volume (GM), total intracranial volume (TIV), and their ratio (GM/TIV). For the subjects that had a measure of cognitive function (i.e. KBIT), we performed multiple regression analyses that included age, sex, and cognitive function as additive predictors of global structural outcome measures (total GM, TIV, and the ratio between the two [GM/TIV]). Two models were conducted per outcome measure, an additive model including the additive effects of the three predictors, and a full saturated model that also estimated their interaction effects. Through stepwise backward regression, we confirmed that the full model was a better fit (AIC/BIC comparison) for GM and GM/TIV but not for TIV. The parameters and estimates for all the additive regression models are presented in Table 6, and the saturated model results are in Table 7. We applied Bonferroni correction for multiple testing comparisons, adjusting for 21 tests (3 outcomes, 7 predictors: 3 main effects, three 2-way interactions, and one 3-way interaction), although this is a stringent correction given the correlations between GM, TIV, and GM/TIV.

**Table 6.** Brain structure and cognitive function linear regression results.

| *Predictors* | **GM** |  |  | **TIV** |  |  | **GM adj TIV** |  |  |
| --- | --- | --- | --- | --- | --- | --- | --- | --- | --- |
|  | *Est* | *CI* | *p* | *Est* | *CI* | *p* | *Est* | *CI* | *p* |
| (Intercept) | -0.39 | -0.51 – -0.27 | **<0.001** | -0.46 | -0.57 – -0.34 | **<0.001** | -0.03 | -0.16 – 0.11 | 0.684 |
| age Year | -0.2 | -0.29 – -0.11 | **<0.001** | -0.15 | -0.24 – -0.06 | **0.001** | -0.15 | -0.25 – 0.05 | 0.005 |
| sex [male] | 0.95 | 0.77 – 1.14 | **<0.001** | 1.11 | 0.93 – 1.29 | **<0.001** | 0.07 | -0.15 – 0.28 | 0.531 |
| KBIT composite | 0.17 | 0.08 – 0.26 | **<0.001** | 0.09 | 0.00 – 0.18 | 0.04 | 0.21 | 0.10 – 0.31 | **<0.001** |
| Observations | 344 |  |  | 344 |  |  | 344 |  |  |
| R<sup>2</sup> / R<sup>2</sup> adjusted | 0.292 / 0.286 |  |  | 0.327 / 0.321 |  |  | 0.107 / 0.088 |  |  |

Est: Estimates; CI: Confidence intervals; p: probability values. P-values in bold indicate statistically significant results.

**Table 7.**
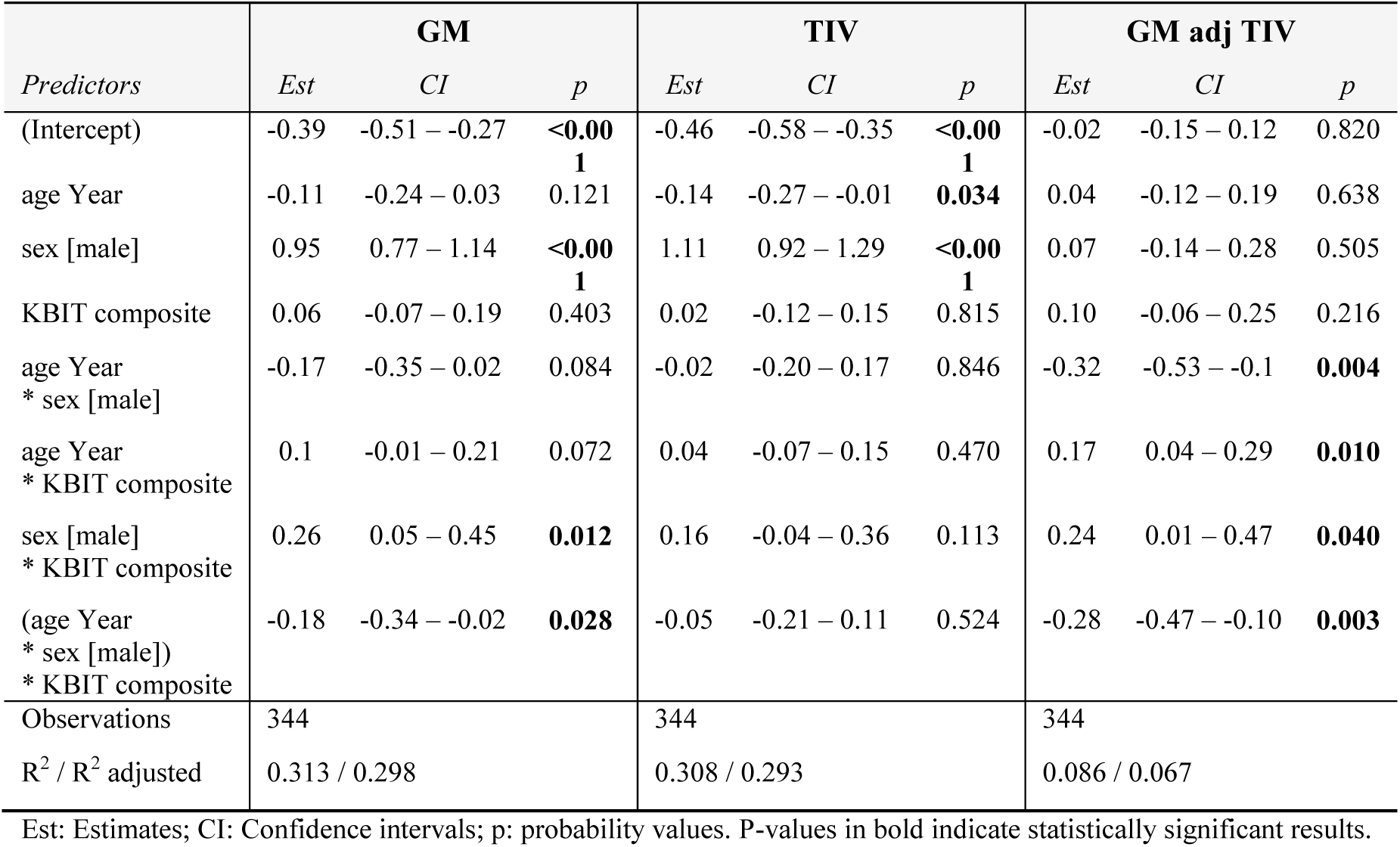
Brain structure and cognitive function linear regression results, full models including interaction terms.

| *Predictors* | **GM** |  |  | **TIV** |  |  | **GM adj TIV** |  |  |
| --- | --- | --- | --- | --- | --- | --- | --- | --- | --- |
|  | *Est* | *CI* | *p* | *Est* | *CI* | *p* | *Est* | *CI* | *p* |
| (Intercept) | -0.39 | -0.51 – -0.27 | **<0.001** | -0.46 | -0.58 – -0.35 | **<0.001** | -0.02 | -0.15 – 0.12 | 0.820 |
| age Year | -0.11 | -0.24 – 0.03 | 0.121 | -0.14 | -0.27 – -0.01 | **0.034** | 0.04 | -0.12 – 0.19 | 0.638 |
| sex [male] | 0.95 | 0.77 – 1.14 | **<0.001** | 1.11 | 0.92 – 1.29 | **<0.001** | 0.07 | -0.14 – 0.28 | 0.505 |
| KBIT composite | 0.06 | -0.07 – 0.19 | 0.403 | 0.02 | -0.12 – 0.15 | 0.815 | 0.10 | -0.06 – 0.25 | 0.216 |
| age Year<br>* sex [male] | -0.17 | -0.35 – 0.02 | 0.084 | -0.02 | -0.20 – 0.17 | 0.846 | -0.32 | -0.53 – -0.1 | **0.004** |
| age Year<br>* KBIT composite | 0.1 | -0.01 – 0.21 | 0.072 | 0.04 | -0.07 – 0.15 | 0.470 | 0.17 | 0.04 – 0.29 | **0.010** |
| sex [male]<br>* KBIT composite | 0.26 | 0.05 – 0.45 | **0.012** | 0.16 | -0.04 – 0.36 | 0.113 | 0.24 | 0.01 – 0.47 | **0.040** |
| (age Year<br>* sex [male])<br>* KBIT composite | -0.18 | -0.34 – -0.02 | **0.028** | -0.05 | -0.21 – 0.11 | 0.524 | -0.28 | -0.47 – -0.10 | **0.003** |
| Observations | 344 |  |  | 344 |  |  | 344 |  |  |
| R<sup>2</sup> / R<sup>2</sup> adjusted | 0.313 / 0.298 |  |  | 0.308 / 0.293 |  |  | 0.086 / 0.067 |  |  |

Est: Estimates; CI: Confidence intervals; p: probability values. P-values in bold indicate statistically significant results.

Sex was consistently associated with GM, TIV, and nominally with GM/TIV, with males having significantly larger volumes across outcomes, while age was negatively associated with GM and TIV but not with GM/TIV. The composite score of cognitive function was positively associated with GM and GM/TIV, but only nominally with TIV.

There was a significant interaction between age and cognitive function predicting GM, and significant age, sex, and cognitive function interactions predicting both GM and GM/TIV (Figure 9). These effects suggest that the relationship between cognitive function and GM (and GM/TIV) is largest in older age, while the association is present in young ages for males but not females. Although these interactive effects captured the variance explained by the main effect of the cognitive function (for GM and GM/TIV) and age (GM), these main effects became non-significant in the full models.

**Figure 9.**
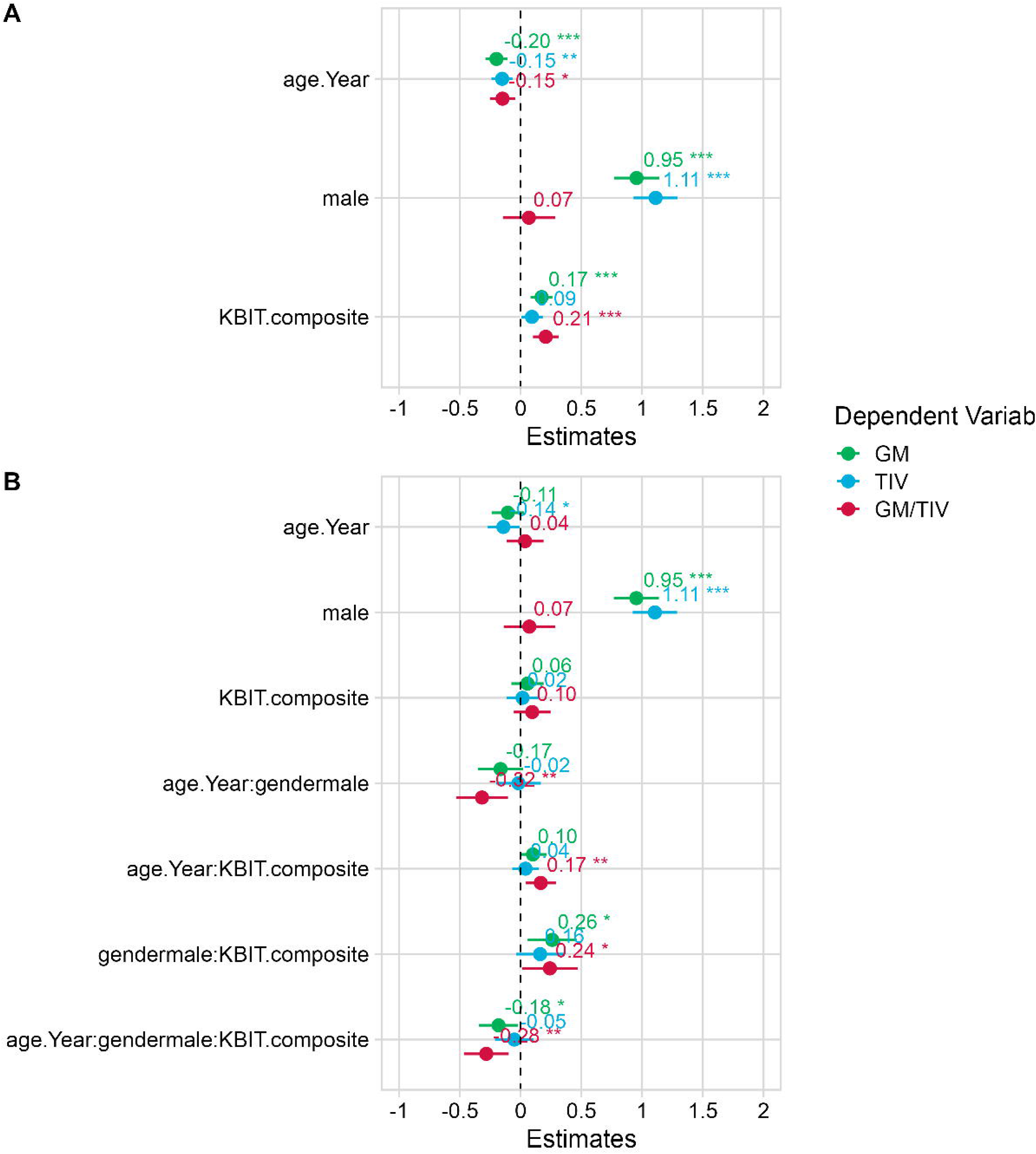
Estimates from structural MRI regressions, including three outcomes: gray matter volume (GM), total intracranial volume (TIV), GM, and their ratio (GM/TIV). A) additive models with age, sex, and cognitive function (KBIT). B) Full models including all interaction terms.

## Usage Notes

The dataset featured in this paper, complete with downloadable materials and codes accessible at <u>NeuroLingua</u> folder on the OpenNeuro platform (https://openneuro.org/), is freely available for use by academic researchers, institutions, and entities, on the condition that proper attribution is given to this work. It is important to note that the authors disclaim any responsibility for the appropriate or inappropriate use in clinical or other contexts. Neither the name of the copyright holder nor the names of its contributors may be used to endorse or promote commercial products derived from this dataset without specific prior written permission.

## Code availability

Custom analysis code relevant to this manuscript can be found in the https://osf.io/b6ur5/.

## Author Contributions

Conceptualization, I.Q., and M.C.; investigation, D.C., P.P.A., C.C., I.Q., A.Sa., L.M.-O.; validation, I.Q., I.D., A.C.-C.; data curation, A.C.-C., I.D., L.D.-S., L.M.-O., M.S., A.Sa., A.Sc., B.C., D.C., and I.Q.; writing—original draft preparation, A.C.-C., I.D., L.M.-O., A.Sa., A.Sc., and I.Q.; visualization, I.Q. and A.C.-C.; formal analysis, I.D., I.Q. and A.C.-C.; funding acquisition, I.Q., A.C.-C., P.P.A., C.C., and M.C. This article is the product of over a decade of coordinated efforts led by M.C. in their role as Scientific Director at BCBL. All authors contributed to various aspects of the research, including data curation, methodology development, manuscript writing, and revisions before submission for review. All authors have read and agreed to the published version of the manuscript.

## Acknowledgments

This research was supported by the Basque Government through the BERC 2022-2025 program and by the Spanish State Research Agency via the BCBL Severo Ochoa excellence accreditation CEX2020-001010-S, as well as the Ramon y Cajal Fellowships RYC2022-035533-I (I.Q.) and RYC-2022-035511-I (A.C.-C.), and the Marie Skłodowska-Curie grant agreement No. 101027016 (A.C.-C.). The authors gratefully acknowledge all the participants who took part in this study.

