## Supplementary information (Figures and Table captions) for "NEUROLINGUA: A Neuroimaging Database Tailored to Unravel the Complexity of Multilingual Comprehension"

S4 Distribution of mean T values across fMRI runs for the top 10 regions of the AAL3 atlas in the VisualPrimary, Auditory and Visual contrasts. The top 10 regions were selected based on the mean T value. For visualization purposes, data were censored to exclude outliers with absolute T-values greater than 30. 4
S5 Visual language comprehension functional topography. Brain renderings and sagittal and axial slices showing probabilistic functional activation for the visual language comprehension > baseline contrast. Activation maps are based on overlaid individual binarized maps, with p-values computed from the same t-maps and combined using Stouffer's method. FDR correction was applied to establish the activation threshold (q = 0.1). This method yielded an alternative atlas that provides a more statistically robust selection of activated voxels. . . 5
S6 Auditory language comprehension functional topography. Brain renderings and sagittal and axial slices showing probabilistic functional activation for the auditory language comprehension > baseline contrast. Activation maps are based on overlaid individual binarized maps, with p-values computed from the same t-maps and combined using Stouffer's method. FDR correction was applied to establish the activation threshold (q = 0.1). This method yielded an alternative atlas that provides a more statistically robust selection of activated voxels. . . . . . . . . . . . . . . . . . . . . . . . . . . . . . . . . . . 6

### Supplementary figures

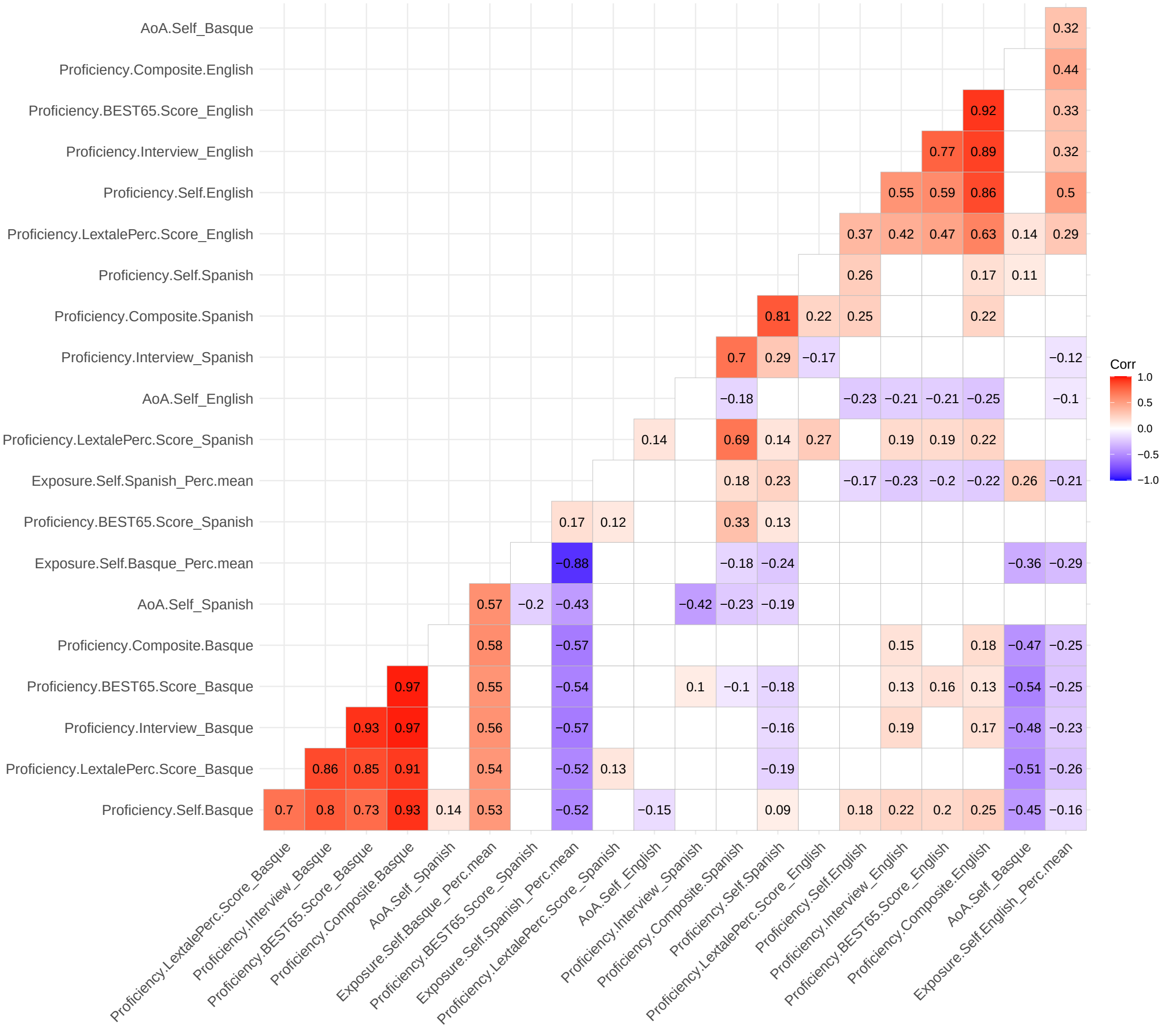

Figure S1: Correlation matrix of linguistic environment variables (Pearson's r). Non-significant values are blanked.

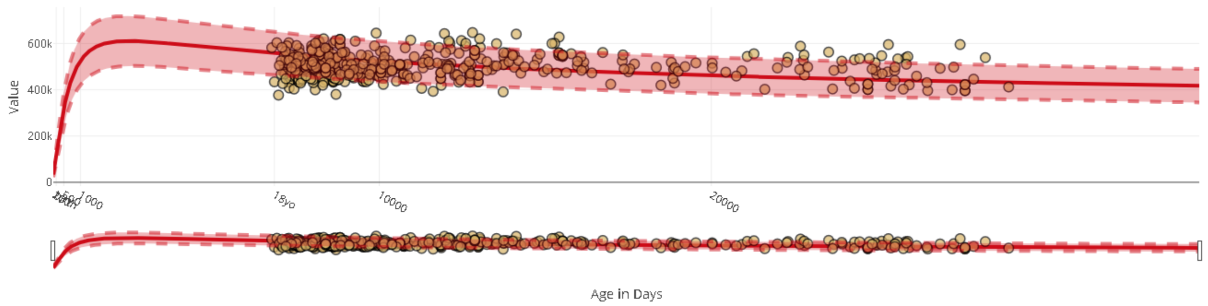

Figure S2: GMV of the NeuroLingua female participants (derived with CAT12) compared to normative braincharts (derived with Freesurfer).

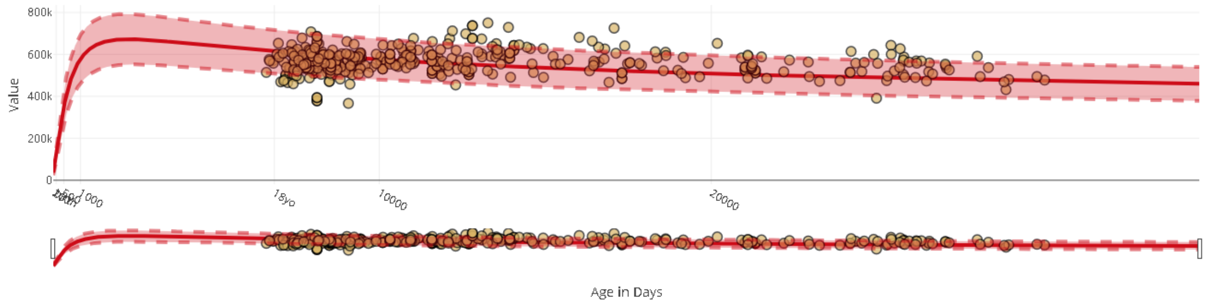

Figure S3: GMV of the NeuroLingua male participants (derived with CAT12) compared to normative braincharts (derived with Freesurfer).

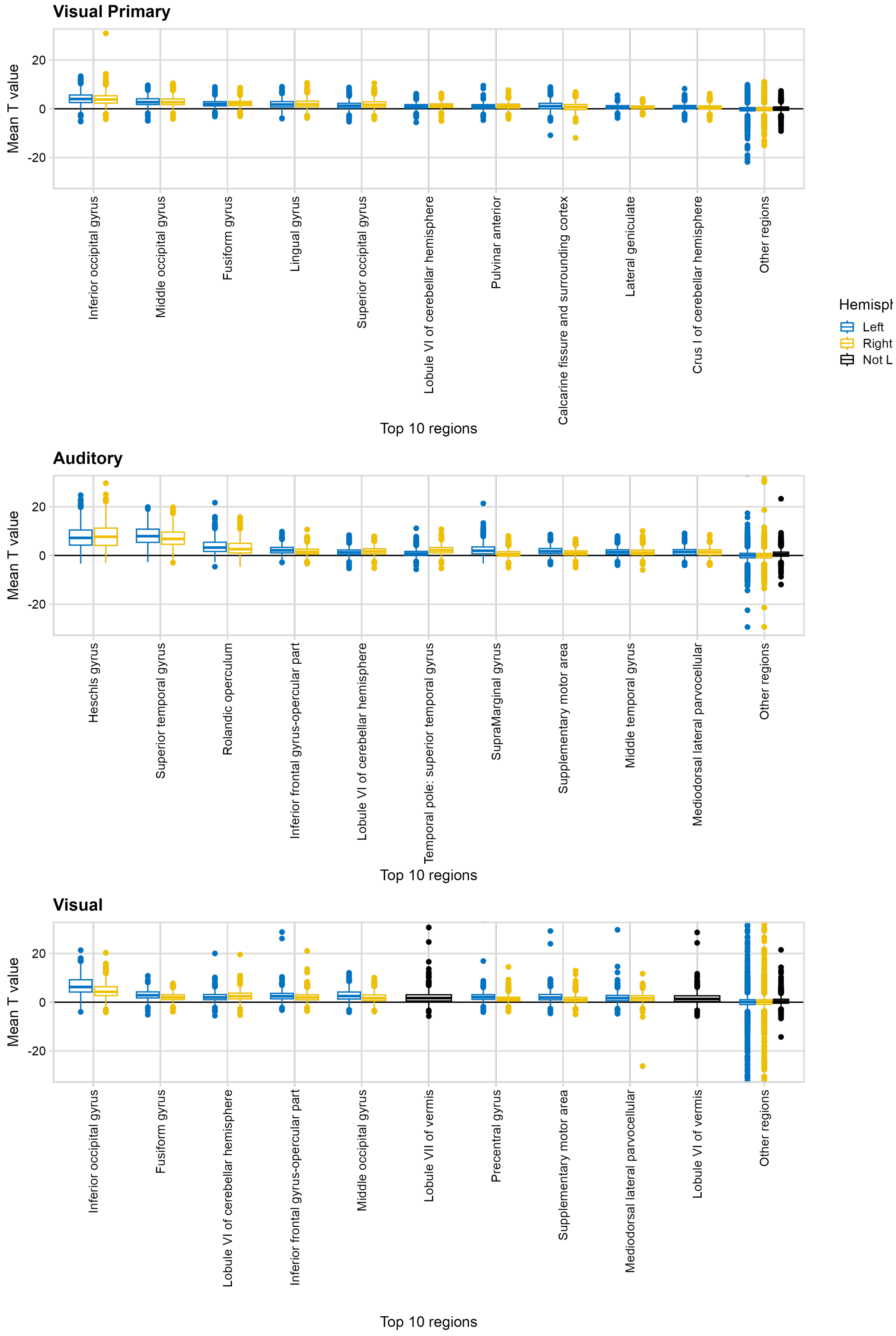

Figure S4: Distribution of mean T values across fMRI runs for the top 10 regions of the AAL3 atlas in the VisualPrimary, Auditory and Visual contrasts. The top 10 regions were selected based on the mean T value. For visualization purposes, data were censored to exclude outliers with absolute T-values greater than 30.

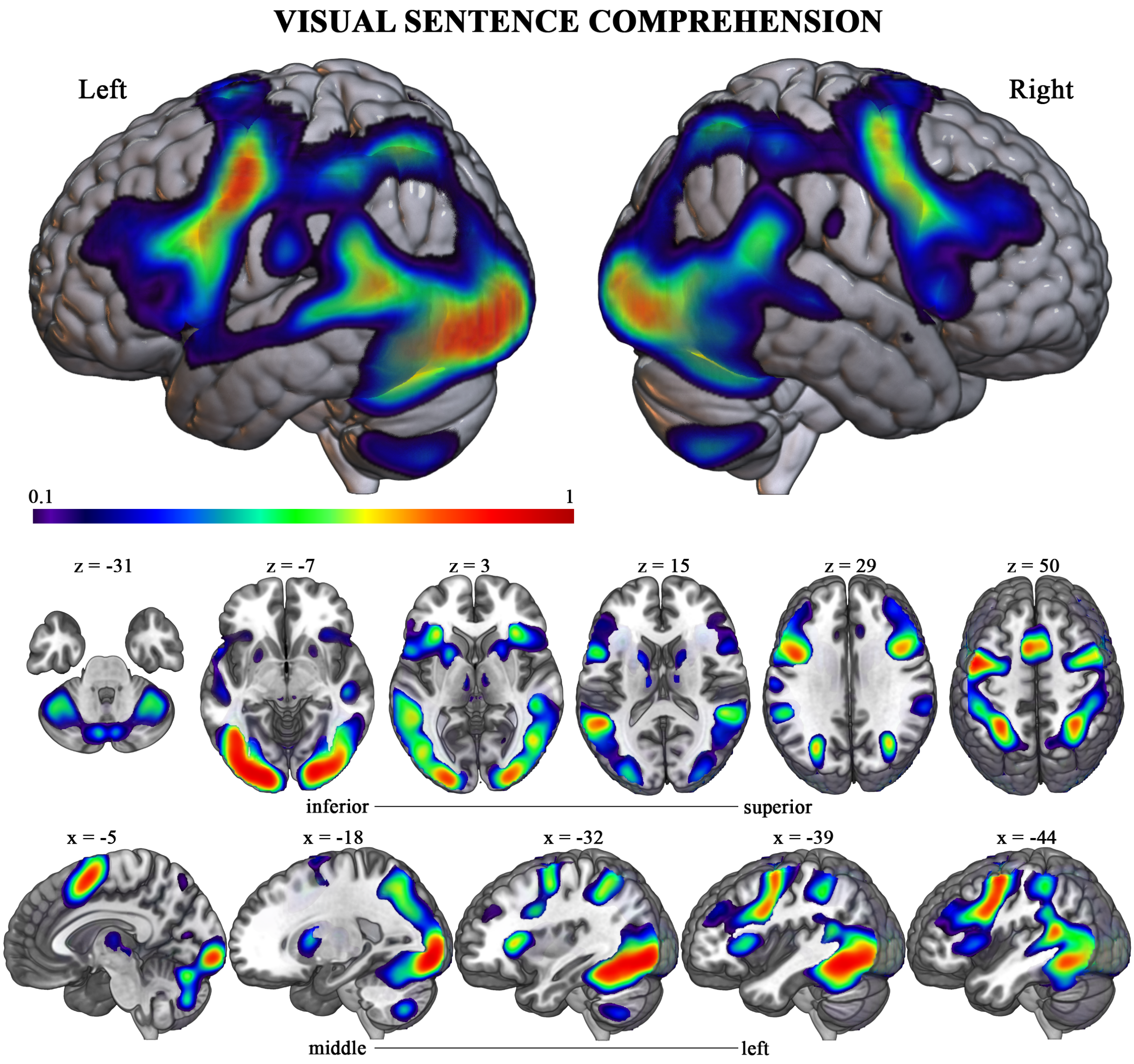

Figure S5: Visual language comprehension functional topography. Brain renderings and sagittal and axial slices showing probabilistic functional activation for the visual language comprehension > baseline contrast. Activation maps are based on overlaid individual binarized maps, with p-values computed from the same t-maps and combined using Stouffer's method. FDR correction was applied to establish the activation threshold (q = 0.1). This method yielded an alternative atlas that provides a more statistically robust selection of activated voxels.

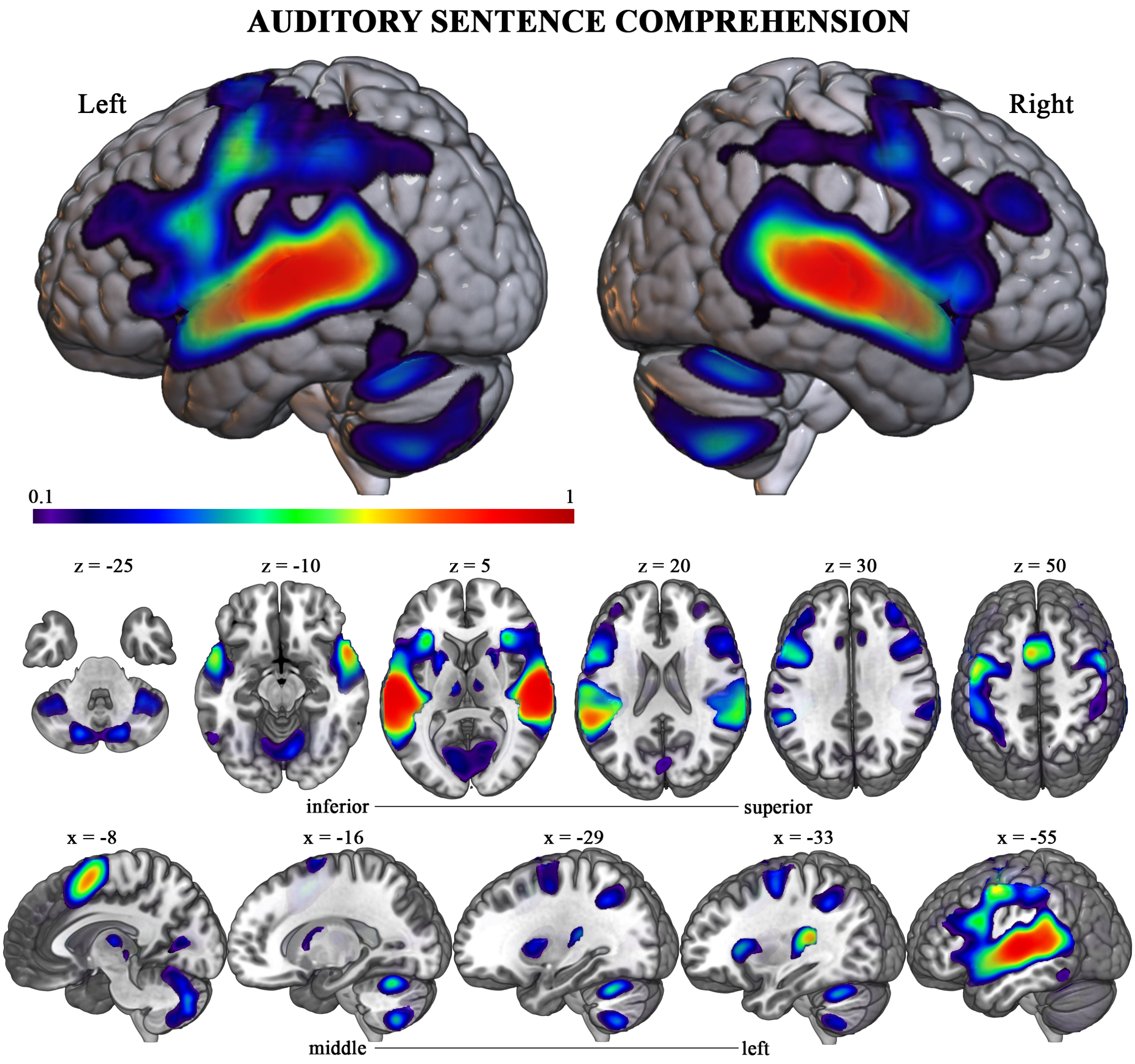

Figure S6: Auditory language comprehension functional topography. Brain renderings and sagittal and axial slices showing probabilistic functional activation for the auditory language comprehension > baseline contrast. Activation maps are based on overlaid individual binarized maps, with p-values computed from the same t-maps and combined using Stouffer's method. FDR correction was applied to establish the activation threshold (q = 0.1). This method yielded an alternative atlas that provides a more statistically robust selection of activated voxels.
